# A Compendium of 49 Experimental SBS Signatures for Decoding Human Cancer Mutational Processes

**DOI:** 10.64898/2026.08.31.748400

**Authors:** Maria Zhivagui, Jessica N Au, Sanskruti Sharma, Peter T Nguyen, Shams Al-Azzam, Jiang Zhang, Mark Barnes, Ludmil B Alexandrov

## Abstract

Human cancer genomes harbor distinct mutational patterns that reflect past processes of DNA damage and repair. However, the precise attribution of these signatures to specific chemical carcinogens lacks a standardized experimental reference framework. To address this gap, we curated 4,282 genome-wide sequencing datasets from 42 model systems across five species exposed to 146 cancer-risk agents. This platform yielded 49 robust experimental single-base substitution signatures (eSS), with 28 matching 19 established COSMIC signatures and 21 defining novel mutational processes. We reconstructed 24 COSMIC signatures, assigning candidate etiologies to five signatures of unknown origin and revising two contested assignments. Pan-cancer decomposition detected four eSS-like mutational processes enriched in smokers across 4,951 tumors. Lastly, independent single-molecule sequencing of primary human organoids reproduced these profiles with high fidelity, confirming true platform-independent biological reproducibility across complex human models. This eSS repertoire provides a reference that links human mutational processes to mechanistic classes of DNA damage.

## Introduction

Somatic mutations accumulate throughout life, recording genotoxic events from conception to disease onset(*1–4*). While many of these mutations arise as a clock-like consequence of endogenous processes, such as the spontaneous deamination of methylated cytosines(*5*), external carcinogenic exposures accelerate this rate and induce unique mutational patterns(*6, 7*). Tobacco smoking accounts for 80-90% of lung cancers(*8*), and UV radiation precipitates 86% of melanomas(*9*). When these environmental insults evade cellular repair machinery, their characteristic DNA lesions become fixed into the genome(*10*). These distinct, recurrent patterns of somatic mutations are termed mutational signatures(*2, 3, 7, 11, 12*). Each signature reflects both the chemistry of the initial DNA lesion and the repair pathways that processed it(*3, 13–15*).

To date, extensive human genomic studies have identified a broad repertoire of mutational signatures associated with specific environmental carcinogens and therapeutic agents, establishing a framework for mutational epidemiology and clinical applications (*2, 12, 16–20*). The COSMIC catalog of mutational signatures features more than 100 single-base substitution (SBS) signatures detected in human cancers, of which a defined subset carries known etiologic assignments: UV radiation (SBS7a/b/c/d), tobacco smoke (SBS4, SBS29, SBS92, SBS100, and SBS109), aristolochic acid I (SBS22a/b/c), aflatoxin B1 (SBS24), colibactin (SBS88), temozolomide (SBS11), and platinum chemotherapy agents (SBS31, SBS35), among others. Nonetheless, many COSMIC signatures retain uncertain or unknown etiology, limiting the utility of mutational signature analysis for causal exposure attribution(*12, 20*).

Therefore, significant efforts have focused on identifying the specific etiologies of cancer mutational signatures(*21*). Controlled experimental models address this challenge by providing a *tabula rasa* genomic background in which direct, causal connections between a defined mutagenic insult and its genomic imprint can be established(*22*). However, a primary technical hurdle lies in accurately distinguishing true somatic mutations from background mutations and sequencing artifacts(*23*). To circumvent this bottleneck, experimental frameworks leverage clonal expansion or tumor formation to amplify true somatic mutations, using platforms that span *in vitro* cell lines and 3D organoids to *in vivo* rodent and nematode models(*22*) (**Figure 1a**).

**Figure 1.**
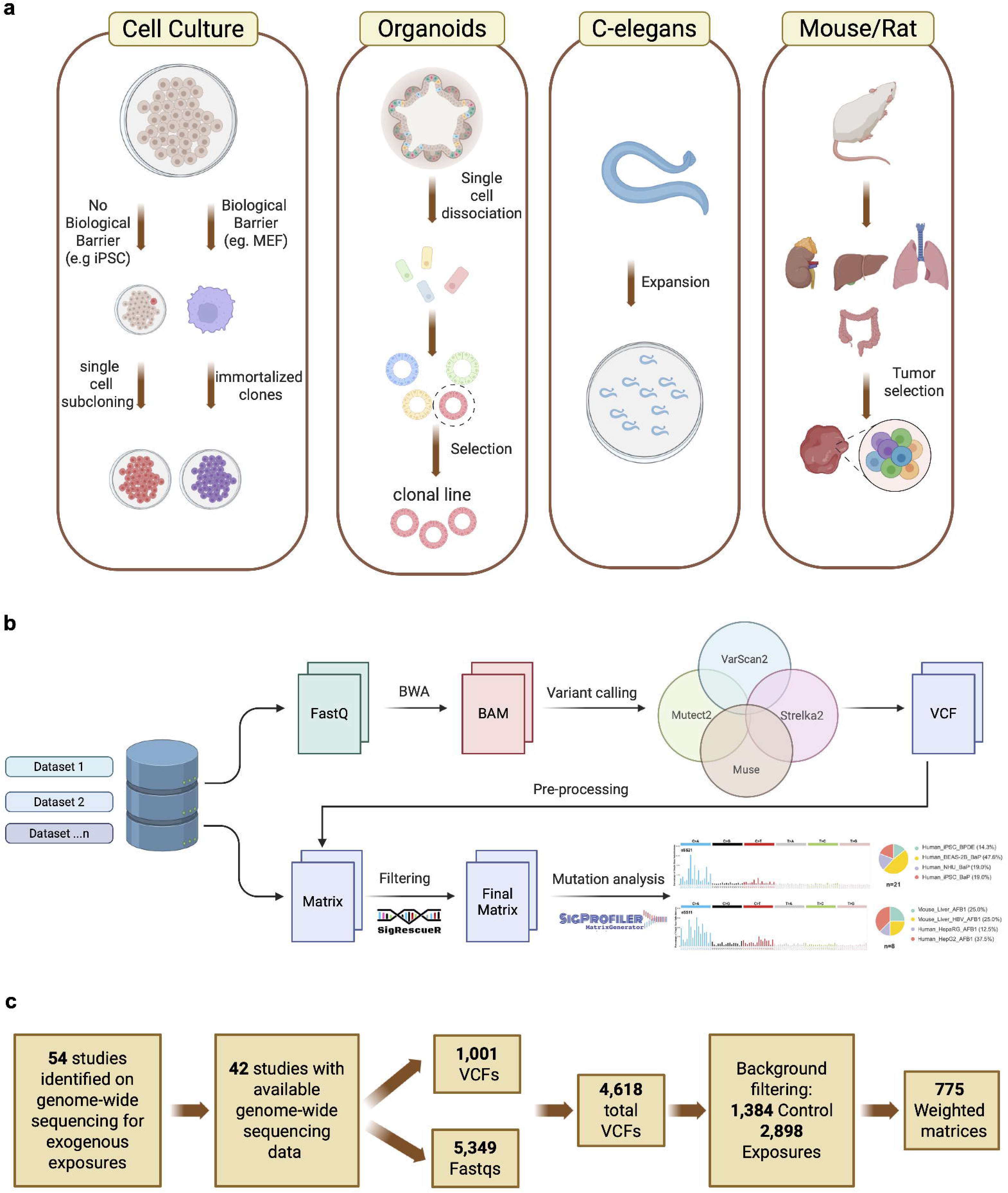
Overview of experimental systems, sequencing pipeline, and data integration. **(a)** Diverse clonal expansion methodologies applied to biological model systems for standard DNA sequencing analysis, featuring cell cultures, 3D organoids, *C. elegans*, and rodent tumor models. In cell culture systems, mutagen exposure was performed followed by single-cell subcloning or serial passaging (3T3 protocol for mouse cells with biological barrier) to generate immortalized clones. Organoid models underwent single-cell dissociation, selection, and clonal line derivation. *C. elegans* experiments involved mutagen exposure and single-cell population expansion. Rodent models included organ-specific exposures and tumor selection before sequencing. (**b)** Depiction of standardized bioinformatics workflow. Raw FASTQ data from multiple datasets were aligned to reference genomes (BWA) to generate BAM files, followed by preprocessing and somatic variant calling (Mutect2, Strelka2, VarScan2, MuSE). High-confidence variants were consolidated into VCF files, filtered, and converted into mutation count matrices. **(c)** Summary of dataset curation and integration. Of 54 studies of genome-wide sequencing for exogenous exposures identified, 42 had available genome-wide data, yielding 4,618 VCFs. Following background filtering, 775 weighted average matrices were generated for downstream analyses.

While more than 50 genome-wide studies have mapped environmental mutagenesis, the field confronts a fragmented landscape. Four fundamental gaps persist: (1) whether carcinogen-induced mutational patterns are reproducible across biological contexts and phylogenetically conserved across species, or are model dependent; (2) which COSMIC mutational signatures are composite signatures that can be deconstructed using a reference set of experimental exposure mutational signatures; (3) how these experimental signatures manifest clinically across pan-cancer cohorts; and (4) which mechanisms of action drive the signatures attributed to environmental exposures.

Here, we compiled 4,282 genome-wide sequencing profiles encompassing 146 cancer-risk agents across 42 experimental systems and five species. From this dataset, we identified 49 reproducible experimental SBS signatures (eSS). These signatures bridge experimental toxicogenomics and human cancer genomics by providing a reference for relating human mutational patterns to mechanistic classes of DNA damage, proposing candidate etiologies for previously unexplained cancer signatures, and defining an evolutionary map of mutagenic responses across metazoans.

## Results

### Framework for Systematic Mutational Profiling Across Model Systems

We first identified 54 studies from which only 42 provide genomic materials (**Figure 1b-c**; **Supplementary Tables 1-2**). We curated a dataset spanning multiple data formats, including FASTQ files, variant calling files (VCFs), and mutation count matrices. Raw sequencing data were reprocessed using a unified analytical framework employing a four-caller consensus strategy to maximize variant detection accuracy and minimize caller-specific artifacts(*24*) (**Figure 1b-c**).

A total of 4,618 VCFs were generated. After pruning samples with incomplete metadata or missing exposure information, 4,282 VCFs remained, encompassing 3,794 whole-genome and 488 whole-exome profiles (**Figure 1b-c**). Somatic mutations were filtered using *SigRescueR* to remove confounding biological and endogenous signals, generating consistent and comparable mutation profiles across all samples, independent of species, model, or experimental conditions(*25*) (**Figure 1b-c**). Samples from the same condition were aggregated into weighted average mutation matrices, enabling downstream comparative analyses across agents and model systems(*25*).

The dataset integrates orthogonal experimental platforms: human-derived models (13.8%), including cancer cell lines, induced pluripotent stem cells (iPSCs), immortalized cells, and 3D organoid models; rodent systems (19.2%), including mouse embryonic fibroblasts (MEFs), tissue-specific mouse organoids, and mouse- and rat-derived tumor tissues; *C. elegans* (66.4%); and the chicken DT40 lymphoblast cell line (0.6%) (**Figure 2a**). *In vivo* tumor-derived models constituted 81% of samples. 88.6% of the datasets were generated by whole-genome sequencing (WGS) (**Figure 2a**). iPSC and *C. elegans* models contributed the largest sample volumes and the most diverse range of tested mutagenic agents (**Figure 2b**; **Suppl. Figure 1**). The number of replicates per model varied widely (median 4; range 1-81; n = 775) (**Suppl. Figure 1-2**). This assembly revealed a marked scarcity of compounds tested across multiple model systems, limiting the high-confidence attribution of mutational signatures across models. 89% of compounds in the compendium were interrogated in one or two model systems, and 78% in only a single species, indicating that cross-species comparisons are achievable for a defined compound subset (**Figure 2c**).

**Figure 2.**
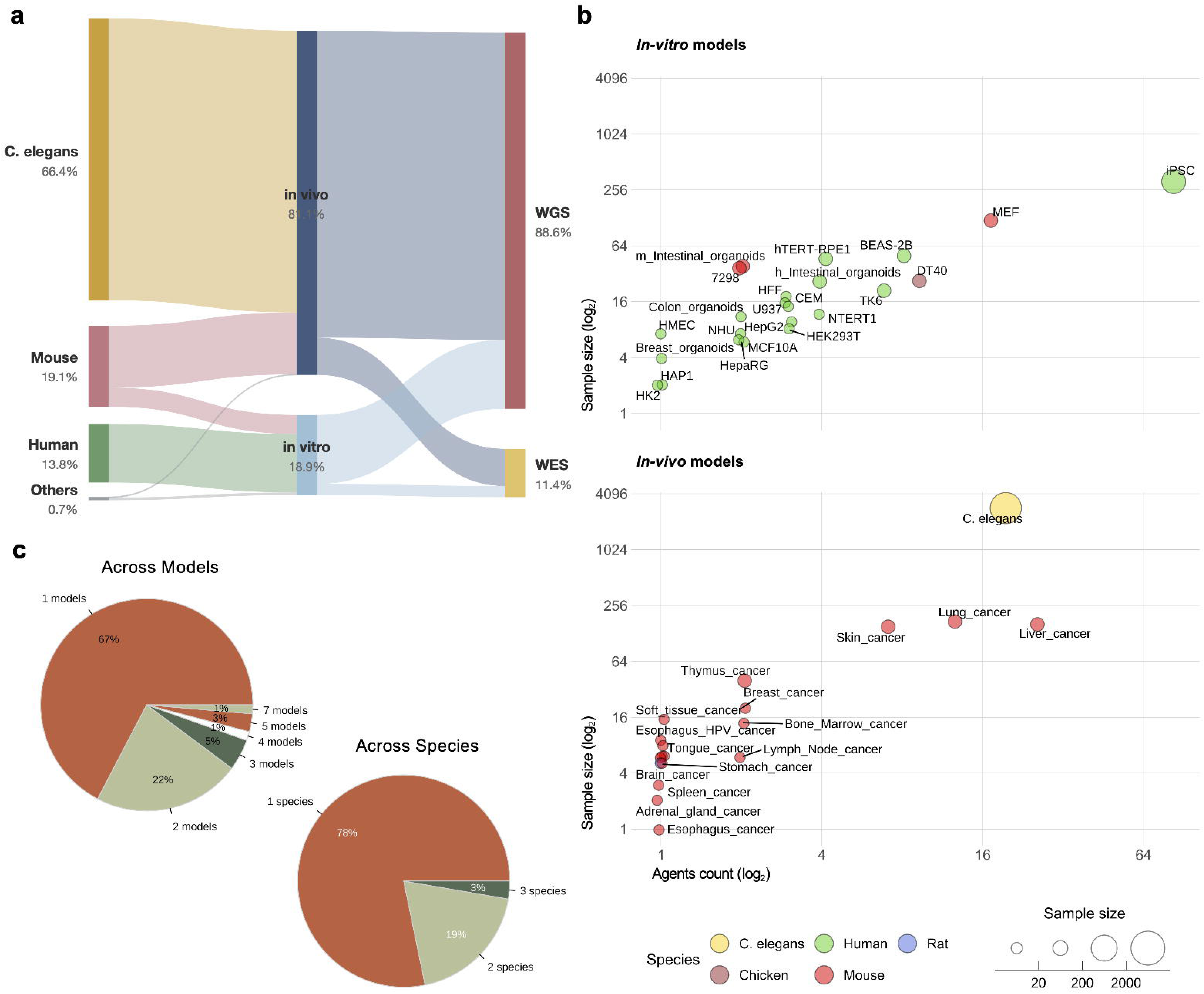
Composition and coverage of the experimental mutational signature atlas. **(a)** Sankey diagram summarizing dataset distribution by species, experimental type, and sequencing strategy. **(b)** Overview of model system diversity and exposure coverage. Top panel shows the *in vitro* models and bottom panel shows the *in vivo* models, represented as the number of distinct mutagenic agents tested (x-axis, log scale) in function of the number of sequenced samples (y-axis, log scale). Point color = species; point size = sample size. **(c)** Percentage of compounds interrogated across models (top panel) and species (bottom panel).

In total, the 4,282 profiles yielded 14,584,637 SBSs. Both filtered and unfiltered mutational landscapes are accessible through the COSMIC experimental Signatures (eSignatures) portal (https://cancer.sanger.ac.uk/signatures/experimental/).

### Mutational Landscapes Across Experimental Systems

We analyzed 1,892,770 SBSs from 554 human genomes and 36 exomes across 19 experimental systems; 4,001,426 SBSs from 364 mouse WGS and 452 WES samples; 53,426 SBSs from 5 rat brain tumors; 8,632,532 SBSs from 2,844 C. elegans WGS samples; and 4,483 SBSs from 27 chicken DT40 WGS samples. To isolate clean, agent-specific mutational patterns with high confidence, background endogenous mutations were systematically removed using the multi-control sample denoising framework in *SigRescueR* and applied independently to each experimental model(*25*) (**Suppl. Figure 3**). Mutational burdens spanned five orders of magnitude across all systems, from hundreds to millions of SBSs, with a subset of treatments exceeding 10 mutations/Mb (**Figure 3a**; **Suppl. Figure 4a**). Ranked by median tumor mutational burden (TMB), the most potent agents across species were methyl methanesulfonate (MMS, 370.2 mut/Mb), pyrrolizidine alkaloids (163.2 mut/Mb), 4-nitroquinoline-1-oxide (4NQO, 79.2 mut/Mb), 5-aza-4-thio-2-deoxycytidine (ATC, 58.1 mut/Mb), solar radiation (29.6 mut/Mb), N-methyl-N-nitro-nitrosoguanidine (MNNG, 24.9 mut/Mb), duocarmycin (21.2 mut/Mb), diethylnitrosamine (DEN, 19.4 mut/Mb), 1,2,3-trichloropropane (TCP, 13 mut/Mb), 9,10-dimethyl-1,2-benzanthracene (DMBA, 12.7 mut/Mb), and trimethyl psoralen with UVA (PUVA, 10.8 mut/Mb). Several compounds, including polycyclic aromatic hydrocarbons (PAHs) such as benzo[a]pyrene (B[a]P) and its diol-epoxide metabolite (BPDE), as well as 4NQO and TCP, induced uniformly high mutation burdens across multiple models, suggesting they target fundamental DNA-chemical affinities that override species-specific differences in repair or metabolism. By contrast, other compounds exhibited model-specific effects. For instance, cisplatin induced high mutation counts in HepG2 cells but not in *C. elegans*, while MNNG was mutagenic in BEAS-2B cells but not in iPSCs. These patterns highlight the influence of model-specific factors, such as metabolic activation and DNA repair capacity, on mutational outcomes (**Figure 3a**; **Suppl. Figure 4a**).

**Figure 3.**
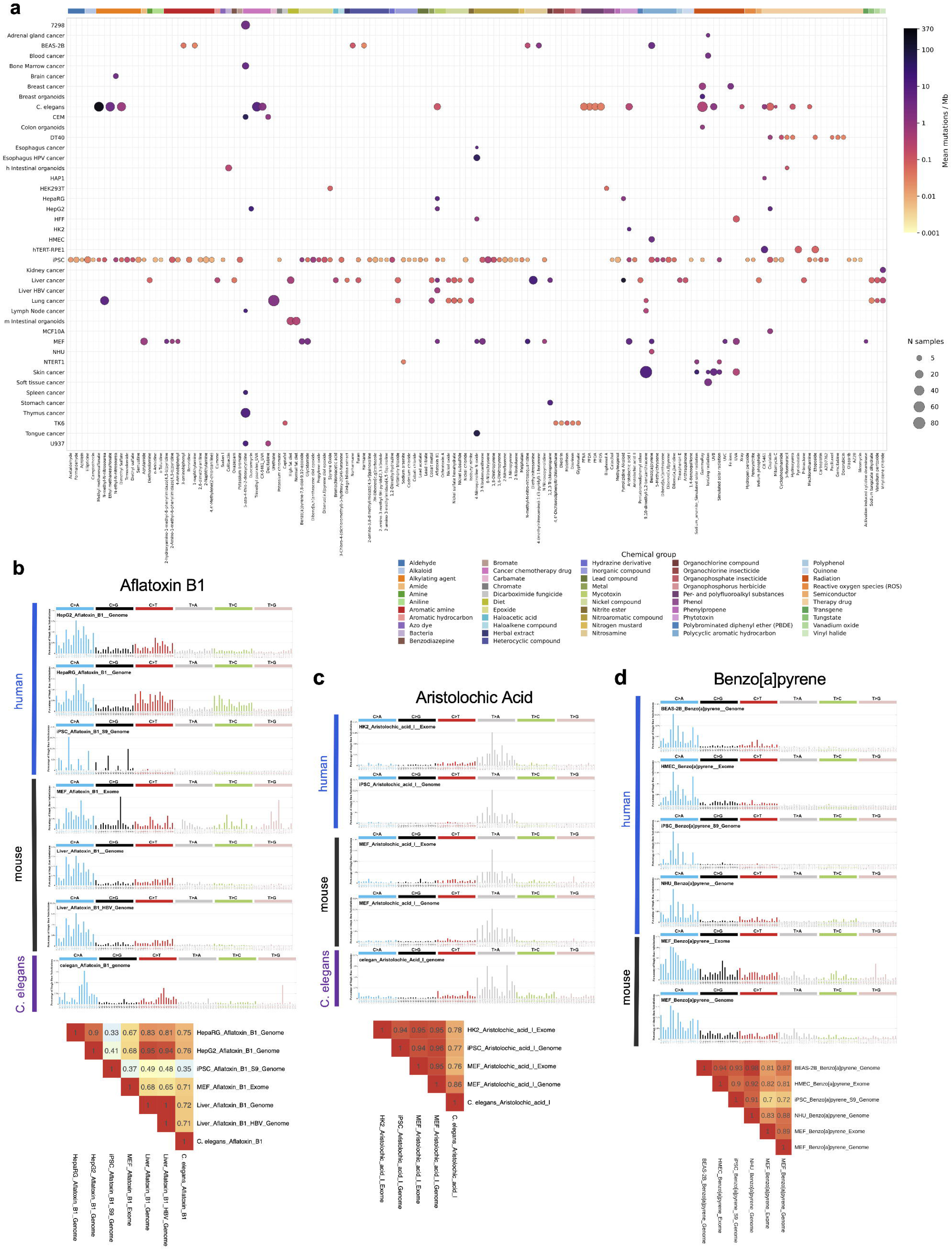
Mutational landscapes and cross-species reproducibility of experimental exposures. **(a)** Tumor mutational burden per mutagenic agent across model systems and species, ranked by median burden (mutation per megabase; log_10_ scale). **(b)** Aflatoxin B1: SBS96 spectra and pairwise cosine similarity across *in vitro* liver models and *in vivo* liver tumors. **(c)** Aristolochic acid I: SBS96 spectra and cosine similarity across species, cell types, and sequencing strategies (WGS and WES). **(d)** Benzo[a]pyrene: SBS96 spectra and cosine similarity across species.

To characterize the global consistency of these chemical insults, we scrutinized 48 compounds that were consistently profiled across species and model systems (**Suppl. Figure 4b-c**). We analyzed the SBS96 mutational patterns of compounds interrogated across more than three distinct model systems (**Figure 3b-d**; **Suppl. Figure 5**). By calculating cosine similarities across diverse experimental conditions, we identified a subset of high-potency agents that leave nearly identical fingerprints regardless of the biological host. HepG2 *in vitro* liver model exhibited mutational profiles that precisely recapitulated the signatures observed in liver cancers from *in vivo* bioassays following aflatoxin B1 exposure (**Figure 3b**), confirming their use as high-fidelity proxies for hepatic-specific carcinogenesis. iPSC-derived models failed to reproduce the characteristic mutational signature seen in the hepatic cells. Aristolochic acid I (AAI) emerged as a remarkably consistent mutagen, producing a nearly identical SBS96 profile across species. This T>A transversion remained biologically constant across sequencing platforms (WGS and WES) and various human and mouse cell types, persisting despite significant differences in DNA repair capacity (**Figure 3c**). Similarly, B[a]P displayed high cross-species stability, dominated by characteristic C>A substitutions, much like solar radiation, which consistently generated C>T mutations across multiple human cell lines and mouse skin tumors (**Figure 3d**; **Suppl. Figure 5e**). In contrast, cisplatin induced highly variable SBS96 profiles, with notable differences between human, chicken, and *C. elegans* models (**Suppl. Figure 5d**). This cross-species map shows that while a mutagen’s SBS96 profile is often conserved across species and cell types, the resulting mutational burden varies dramatically by cellular context (**Suppl. Figure 5**).

### A High-Resolution Compendium of 49 Experimental SBS Signatures

To ensure signature fidelity, we evaluated mutational profile stability using Poisson resampling (see STAR methods)(*26*). Evaluating 1,000 Poisson resamples across 4,282 mutational profiles showed that low-mutation profiles fluctuate widely, whereas high-mutation profiles remain virtually unchanged (**Suppl. Figure 6a**). These samples required a minimum of 307 SBSs to withstand random error, above which 95% of resampled profiles achieved a cosine similarity ≥0.95 with the original (**Suppl. Figure 6a**). Profiles exceeding this threshold accounted for 15.7% of the samples (671 out of 4,282) across 74 unique exposures (**Suppl. Figure 6b**).

Hierarchical clustering of the 671 high-confidence profiles identified 49 distinct eSS that defined the somatic landscape across tissues and species with ≥3 samples per cluster (**Figure 4a**; **Suppl. Figure 7a**; **Supplementary Tables 3-4**). The 49 eSS represent a conservative, high-confidence subset of the detectable mutagenic signal across the compendium of 50 exposure treatments from 508 samples. Certain agents exhibited sample-level mutational stochasticity, that is, heterogeneity among replicates within the same agent, that precluded clustering into a consensus signature (**Suppl. Figure 7b**; **Suppl. Figure 8**). Profiles that did not generate profiles meeting the ≥3-sample reproducibility criterion required for eSS cluster assignment were extracted as small clusters (n = 2) and singletons (n = 1), accounting for 163 samples across 49 unique compounds (**Suppl. Figure 7b; Suppl. Figure 8**; **Supplementary Tables 5-6**). In human cellular models, pyridostatin generated substantial mutation burdens but produced highly heterogeneous replicate profiles that appeared as singletons (**Suppl. Figure 8**). Similarly, *C. elegans* strains exposed to gamma radiation imprinted high TMB compared to the non-treated samples and failed to cluster together.

**Figure 4.**
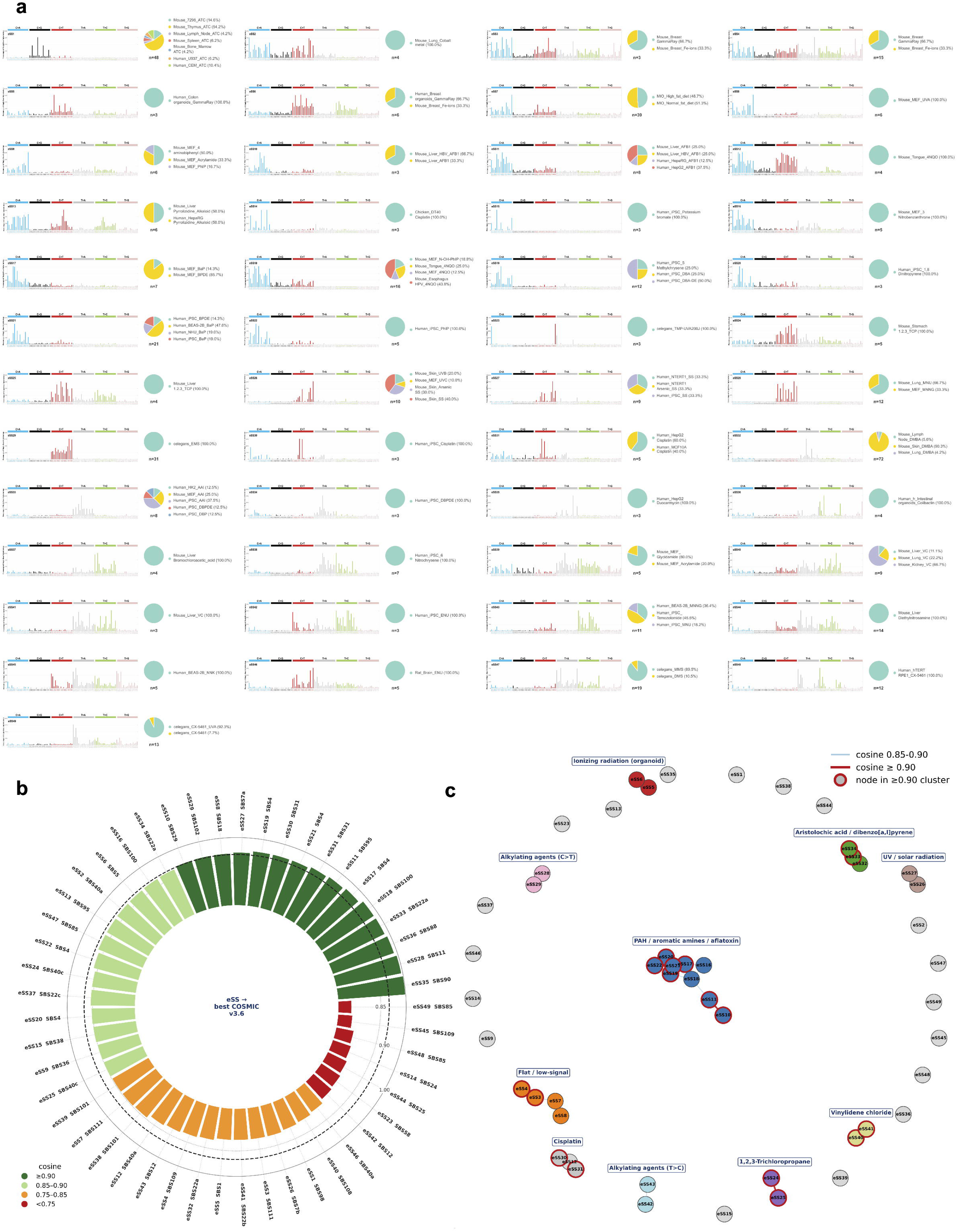
A compendium of 49 experimental SBS signatures and their pattern relationships. **(a)** Hierarchical clustering of 671 high-confidence profiles (≥ 307 substitutions) identified 49 experimental SBS signatures (eSS), each supported by ≥ 3 samples. For each eSS, the SBS96 profile is shown alongside a pie chart illustrating sample contributors, including their relative percentages, total counts, experimental models, and compound treatments. **(b)** Cosine similarity between the 49 eSS and COSMICv3.6 SBS signatures. **(c)** Pattern family structure of the 49 eSS, partitioned into seven families defined by dominant mutation type and chemical class.

To benchmark our experimental models against human data, we compared the 49 eSS against the COSMICv3.6 database. More than half of our signatures (28 eSS; 57%) demonstrated high concordance (cosine similarity ≥ 0.85) with 19 established COSMIC signatures, successfully capturing canonical mutational processes(*27*). 15 of these eSS exhibited exceptional alignment (≥ 0.90) with known signatures, yielding highly precise recapitulations, such as eSS28 matching the alkylating signature SBS11 (0.99) and eSS35 perfectly mirroring SBS90 (1.00) (**Figure 4b**; **Suppl. Figure 9**; **Supplementary Table 7**). Intriguingly, the remaining 21 eSS lacked any COSMIC counterparts. These *de novo* signatures represent previously uncharacterized mutational processes, expanding the repertoire of experimentally defined mutational processes.

To assess whether these compounds generated detectable mutagenic signal despite failing the reproducibility threshold, we compared the singleton clusters and small clusters of n = 2 against the full COSMICv3.6 SBS database and the 49 eSS using cosine similarity, testing for any signature missed by the 49-eSS compendium (**Suppl. Figure 10**). Three clusters reached cosine similarity ≥ 0.9 to at least one COSMICv3.6 signature while remaining absent from the eSS compendium (**Suppl. Figure 10**). These clusters generated mutational signal that aligned with established cancer processes despite insufficient replication for independent eSS derivation. SBS7c matched both UVA exposure in human dermal fibroblasts (HFF; cosine similarity = 0.905) and the alkylating agents, ethyl methanesulfonate (EMS) and dimethyl sulfate (DMS), in *C. elegans* (cosine similarity = 0.92). This convergence reflects shared thymine-directed chemistry. EMS and DMS alkylate thymine at the O^2^ and O^4^ positions(*28*), while UVA generates thymine-containing cyclobutane pyrimidine dimers, both directly and through reactive oxygen intermediates(*29*). Next, 5-fluorouracil (5-FU) matched SBS17b (cosine similarity = 0.96), corroborating reported clinical associations between 5-FU chemotherapy and SBS17(*30, 31*). This alignment demonstrates that exogenous uracil analogs and endogenous dNTP pool oxidation(*32*) converge on a shared mutational spectrum (**Suppl. Figure 11**).

The remaining clusters showed no confident COSMIC match and represent the true experimental negatives of the compendium: agents that either did not generate detectable above-background point mutations in the tested systems, or whose signal was too weak and heterogeneous to resolve against the COSMIC reference.

### Pattern relationships of the 49 experimental signatures

We compared the 49 eSS by cosine similarity and grouped those scoring ≥ 0.90, yielding 10 clusters. Each cluster is defined by a dominant mutation class and chemical mechanism (**Figure 4c**; **Suppl. Figure 12**). The PAH and nitroaromatic family is C>A-dominated (54 to 77% C>A) and the largest coherent module. It contains the B[a]P/BPDE signatures eSS17 (mouse) and eSS21 (human; mutual cosine 0.907), eSS16 (3-nitrobenzanthrone), eSS18 (4NQO and N-OH-PhIP), eSS19 (5-methylchrysene), eSS20 (1,8-dinitropyrene), and eSS22 (PhIP). All match SBS4 or SBS100 at cosine similarity between 0.86 to 0.97 (**Suppl. Figure 9**). Within the C>A module, the aflatoxin B1 signatures eSS10 and eSS11 (mutual cosine similarity = 0.90) form a peripheral sub-cluster connected to the PAH/nitroaromatic core by a single link (eSS11 vs. eSS18, cosine similarity = 0.89; **Figure 4c**), while remaining otherwise distinct (cosine similarity ≤ 0.84 to all PAH members). At a stricter threshold (cosine similarity ≥ 0.90), the aflatoxin pair separates entirely (**Suppl. Figure 12**). This isolation reflects a mechanistically distinct C>A process driven by bulky N^7^-guanine adducts that overlaps with the PAH family in pattern yet remains structurally separable.

Alkylating agents did not form a single mutational module but split into three substitution classes defined by the alkylated atom (**Figure 4c**). Two C>T-dominated signatures, eSS28 (mouse; N-methyl-N-nitrosourea, MNU, and MNNG, 92% C>T) and eSS29 (EMS, 89% C>T), clustered together (mutual cosine similarity = 0.86) and matched SBS11 (cosine 0.99) and SBS102 (cosine similarity = 0.91), respectively. These signatures reflect an O^6^-alkylguanine process in which the alkylated guanine mispairs with thymine(*33, 34*). Two T>C-dominated signatures, eSS42 (N-ethyl-N-nitrosourea, ENU, 53% T>C) and eSS43 (human; MNU, MNNG, and temozolomide, 65% T>C), clustered separately (mutual cosine similarity = 0.88). Both can arise from O^4^-alkylthymine which mispairs with guanine to produce T>C transitions(*33*). O^4^-alkylthymine is poorly repaired and concentrates mutations at A:T base pairs, whereas the co-produced O^6^-alkylguanine, the source of C>T, is likely removed by O^6^-methylguanine-DNA methyltransferase (MGMT) (*33, 34*). This repair asymmetry likely accounts for the divergence between eSS28 and eSS43: the same methylating agents generated a C>T signature in rodent tissue but a T>C signature in human iPSC. The C>T and T>C clusters shared almost no similarity (cosine similarity ≤ 0.27), which places them in distinct mutational processes despite a common alkylating origin. Two further signatures remained singletons at this stringency: eSS47 (MMS and DMS) is T>A-dominated and matched SBS85 (cosine similarity 0.88) and eSS46 (ENU in rat) is a mixed profile of low similarity to any reference (**Figure 4c**).

The T>A family comprises three signatures that peak at C[T>A]G: AAI (eSS33; cosine similarity to SBS22a = 0.97), dibenzo[a,l]pyrene and its diol epoxide metabolite (DBP and DBPDE; eSS34; cosine similarity to SBS22a= 0.89), and DMBA (eSS32; cosine similarity to SBS22a = 0.80). The three signatures exhibit 60 to 73% T>A mutation class (**Figure 4c**). The module spans human and mouse exposures yet retains high similarity (cross-species cosine similarity between 0.87 to 0.92), making it one of the most reproducible chemically defined families (**Figure 3c**; **Figure 4c**).

The UV/solar radiation cluster reflects a cross-species C>T pair at dipyrimidine contexts: eSS26 (mouse skin; 76% C>T) and eSS27 (human keratinocyte and iPSC; 74% C>T). Cisplatin formed a defined C>T cluster (eSS30, eSS31) matching SBS31 at cosine 0.92 to 0.94. Conversely, cisplatin exposure in chicken cells (eSS14, DT40) produced a C>A-enriched pattern (64% C>A) and remained a singleton, consistent with a different repair background processing the same platinum adducts(*35*) (**Figure 4c**).

Finally, four smaller clusters resolved outside the major etiologic modules. TCP formed a separate C>T cluster (eSS24, eSS25). Vinylidene chloride formed its own cluster that split by tissue, eSS40 (kidney, lung, and liver) and eSS41 (liver), both with mixed T>C and T>A. Ionizing radiation from gamma rays and Fe-ions split across two low-similarity clusters, one in organoids (eSS5, eSS6) and one in *p53*-mutant mammary tumors that also drew in C>A signals from fat diet organoids and UVA (eSS3, eSS4, eSS7, eSS8), neither forming a chemically defined module (**Figure 4c**).

### Phylogenetic Conservation of Experimental Signatures Across Species

More than half of the identified clusters (26/49) comprised a single sample type, reflecting high compound-specificity and technical robustness. The remaining 23 clusters demonstrated stability across multiple compounds, models, or species: 65.2% (15/23) spanned multiple models and 69.6% (16/23) involved multiple compounds. Cross-species clustering, though less common (21.7%), consistently identified high-potency carcinogens (**Figure 4c**; **Supplementary Table 8**).

For compounds with multi-species coverage, we evaluated the pairwise cosine similarities between eSS derived from the same agent across different model organisms (**Figure 5a**). Pairwise cosine similarities varied substantially between signatures, ranging from 0.181 to 0.907, highlighting a distinct mechanistic division between agents whose signatures are strictly chemically determined and those dictated by host repair context. This data resolved into a clear hierarchy where high cross-species conservation is restricted to agents that produce highly reproducible, chemically defined DNA lesions (**Figure 5a**). For instance, B[a]P/BPDE (human vs. mouse: 0.907) and solar simulation (human vs. mouse: 0.897) are strongly conserved. Both produce lesions, specifically bulky guanine adducts and dipyrimidine photoproducts, respectively, whose sequence-context specificity is dictated by the intrinsic chemistry of the DNA-mutagen interaction rather than by the biological host.

**Figure 5.**
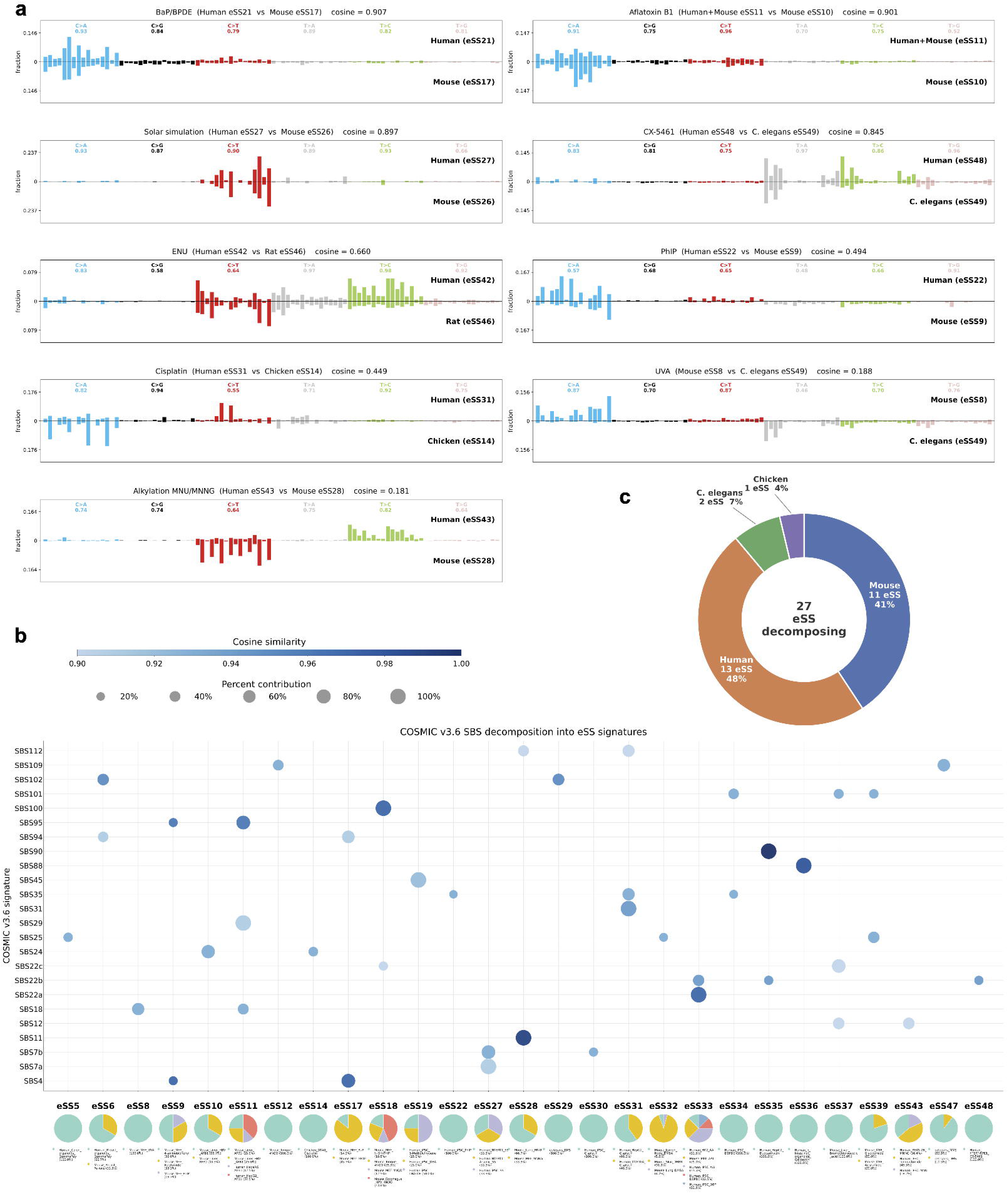
Cross-species conservation and decomposition of COSMICv3.6 signatures by the eSS compendium. **(a)** Pairwise cosine similarity between eSS derived from the same agent across model organisms. **(b)** Decomposition of COSMICv3.6 SBS signatures against the 49 eSS (SigProfilerAssignment, reconstruction cosine ≥ 0.90). Point size denotes the percent contribution of each eSS; color denotes cosine similarity. **(c)** Pie chart of the origin of the reconstructing eSS per species.

Conversely, alkylating agents displayed pronounced inter-species divergence: ENU showed weak conservation (human vs. rat: 0.660), while MNU and MNNG exhibited near-complete divergence (mouse vs. human: 0.181). Despite sharing an alkylation mechanism that produces a characteristic O^6^-alkylguanine-driven C>T pattern, this footprint is heavily modulated by species-specific differences in MGMT repair efficiency(*33, 36*). Low conservation was similarly observed for cisplatin (chicken vs. human: 0.449), PhIP (mouse vs. human: 0.494), and UVA (mouse vs. *C. elegans*: 0.188; **Figure 5a**). Thus, signature identities are largely dictated by host repair context, including the variable complement and kinetics of NER, BER, and MMR, rather than DNA damage chemistry alone.

### Experimental SBS Signatures Resolve Rare COSMICv3.6 Signatures

To establish a mechanistic bridge between controlled experimental exposures and the complex mutational patterns observed in human clinical cohorts, we decomposed COSMICv3.6 signatures against the compendium of 49 eSS using SigProfilerAssignment, retaining reconstructions with cosine similarity ≥0.90 (**Figure 5b**; **Supplementary Table 9**). This framework revealed that 27 distinct eSS successfully reconstructed 24 COSMIC SBS signatures, originating predominantly from human (48%) and mouse (41%) models (**Figure 5c**). Notably, a high reconstruction cosine similarity establishes pattern compatibility and not a unique solution. Because several eSS are mutually correlated (**Figure 4c**), a given COSMIC signature can admit more than one eSS combination (**Supplementary Table 10**). Identifiability testing partitioned the 24 reconstructed COSMIC signatures into 11 robust reconstructions and 14 reconstructions with ambiguous solutions (**Supplementary Table 10**). We therefore treat each reconstruction as a mechanistic decomposition rather than a specific agent attribution. Bootstrap resampling confirmed that the eSS solution of every reconstruction cited below was selected in ≥ 95% of 1,000 resamples, while components below 5% weight were unstable across resamples and are therefore not reported.

Crucially, COSMIC decomposition recovered established clinical etiologies (**Figure 5b**). For example, SBS4, the signature of tobacco-associated lung and laryngeal cancers, was reconstructed as a multi-component mixture of PAHs and related mutagens, comprising 75.7% eSS17 (B[a]P/BPDE in MEFs) and 24.3% eSS9 (PhIP and acrylamide in MEFs)(*37*). Although SBS4 was flagged as ambiguous due to degenerate solutions, the qualifying alternative eSS combinations were strictly constrained to the same chemical families. Across all qualifying alternatives, six of the nine contributing eSS belonged to the PAH class and three to aromatic or heterocyclic amines. This chemical class stability confirms that while individual agent attributions remain interchangeable, the multi-component assignment reflects the genuine, multi-class chemical complexity of tobacco smoke mutagenesis rather than mathematical instability(*37, 38*).

Among the robust reconstructions, SBS7a decomposed entirely into eSS27 (UV exposure in human cells, 100%), while SBS7b assigned 74% of its density to eSS27, yielding high cosine similarities of 0.91 and 0.93, respectively. The alkylating agent-associated signature, SBS11, mapped entirely to eSS28 (MNU and MNNG in MEFs) with an exceptional cosine similarity of 0.99. Similarly, the AAI exposure-associated mutational signature SBS22a decomposed completely into eSS33 (AAI across human HK-2, iPSC, and MEF models), while its clinical variant SBS22b assigned 47% to eSS33 (cosine similarity = 0.94). The platinum-induced signature SBS35 was reconstructed using eSS31 (cisplatin in human HepG2 and MCF10A cells), which contributed 58% of its profile with a cosine similarity of 0.93. For niche environmental exposures, the colibactin-related signature SBS88(*39, 40*) was accurately reconstructed from eSS36 (colibactin in human intestinal organoids, 100%) with a cosine similarity of 0.98, and SBS90 mapped entirely to eSS35 (duocarmycin exposure in HepG2) with a cosine similarity of 0.998 (**Figure 5b**).

In contrast, for signatures admitting multiple equivalent eSS combinations, individual agent attributions were ambiguous, but chemical class etiologies remained consistent. The platinum-induced signature SBS31 was successfully reconstructed using eSS31 (cisplatin), which contributed 100% of its profile with a cosine similarity of 0.935. Additionally, the tobacco smoke-associated signature SBS100 was fully captured by eSS18 (4NQO in mouse models) with a cosine similarity of 0.97, and the aflatoxin B1 signature SBS24 reconstructed from the mouse liver aflatoxin B1 signature eSS10 (70.2% assignment, reconstruction cosine similarity 0.93). Finally, the clinical variant SBS22c was assigned 74% to eSS37 (bromochloroacetic acid in liver; cosine similarity = 0.903). (**Figure 5b**; **Suppl. Figure 13a**).

Beyond these established assignments, our experimental framework resolved seven COSMIC signatures of unknown or contested etiology (**Figure 5b**). Of these, SBS29 and SBS109 possessed only speculative clinical annotations, for which the compendium proposes candidate experimental etiologies. The remaining five signatures (SBS94, SBS95, SBS101, SBS102, and SBS112) lacked any prior etiological framework and received putative experimental assignments.

Although clinically annotated as a tobacco-chewing signature, SBS29 is reconstructed here entirely by the human aflatoxin signature eSS11 (100% assignment, cosine similarity = 0.91). When extracted from human cells exposed to aflatoxin B1, eSS11 matches SBS29 more closely than it matches SBS24, yielding a similarity of 0.91 versus 0.88, and carries an excess of A[C>A]A and C[C>A]N mutations, consistent with a shift in the aflatoxin mutational spectrum in the human epithelial context (**Suppl. Figure 13b**). This attribution links SBS29 to an aflatoxin-like mutational process in lung tissues, which likely represents a mathematical artifact driven by C>A spectral overlap with tobacco smoke rather than genuine aflatoxin exposure. (**Suppl. Figure 13b**).

SBS94 reconstructs from eSS6 and eSS17 at cosine similarity 0.91, with the B[a]P experimental signature, eSS17, as the dominant component (62%) (**Figure 5b**; **Suppl. Figure 14a**). SBS95, previously reported as an artefactual liver signature(*41*), reconstructs primarily from eSS11 at 76% with a cosine similarity of 0.96 (**Suppl. Figure 14b**). The human aflatoxin experimental signature, eSS11, matches SBS95 with a similarity of 0.94, representing the strongest single eSS match to this signature in the entire compendium (**Suppl. Figure 9**). Consequently, SBS95 groups with the dietary hepatotoxin family, linking some of its clinical presentation to aflatoxin or related mycotoxin exposure (**Suppl. Figure 14b**).

The recently described liver signature, SBS101, robustly reconstructs from three nearly equal contributors to achieve a cosine similarity of 0.93 (**Figure 5b**): eSS34 (38%), eSS37 (33%), and eSS39 (29%). Moreover, SBS102 reconstructed from an almost equal split between eSS29 (EMS) and eSS6 with a cosine similarity of 0.95 (**Figure 5b**). This specific composition assigns SBS102 to an active ethylation process operating on a background of human aging.

As a signature associated with tobacco smoking, SBS109 reconstructs from eSS47 and eSS12 with a cosine similarity of 0.93. The methylating agents’ signature eSS47 (MMS and DMS) is dominant at 60% (**Figure 5b**). eSS12 reproduces the C>A and C>T arms of SBS109 at per-class cosine 0.88 and 0.91, respectively. eSS47 reproduces the T>A and T>C arms at per-class cosine 0.94 and 0.89, respectively. This alkylation origin strongly implicates tobacco-derived alkylating carcinogens in head, neck, and lung tissues(*42, 43*).

Finally, SBS112, a rare signature of unknown cause, reconstructs from the alkylating signature eSS28 and the platinum signature eSS31 with a cosine similarity of 0.90 (**Figure 5b**). The platinum signature eSS31 supplies the dominant 56% and drives the primary C>T signal, while the alkylation profile eSS28 contributes 44%.

### Per-class mutation similarity between eSS and COSMICv3.6 signatures reveal plausible therapeutic associations

Signature eSS1 is derived exclusively from exposure to the cytidine analog ATC, demonstrating a highly conserved mutational pattern across multiple species and experimental models. This signature is defined by a pronounced C>G enrichment concentrated in three distinct N[C>G]G trinucleotide contexts, specifically C[C>G]G at 23.4%, A[C>G]G at 14.0%, and G[C>G]G at 14.0%, which collectively account for 51.4% of the total profile(*44*). The full 96-channel profile of eSS1 showed partial similarity to COSMICv3.6 SBS98 with a cosine similarity of 0.78.

Decomposition of SBS98 against the eSS anthology resolved the clinical signature into eSS1 at 47.6%, achieving a reconstruction cosine similarity of 0.845 (**Suppl. Figure 15a**). Scrutinizing individual mutation classes revealed that the C>G and C>A patterns of COSMIC SBS98 both share a high cosine similarity of 0.93 for both with eSS1 (**Suppl. Figure 15b**). This structural alignment associates clinical SBS98 with cytidine-analog chemotherapy exposure. Cytidine-analog DNA methyltransferase inhibitors are used to treat leukemia, and SBS98 is detected in acute lymphoblastic leukemia, the lineage in which ATC drove C>G mutagenesis and leukemic transformation in mice(*44*). Because it lacks a high-confidence COSMIC match, eSS1 is retained as a unique novel signature, and its partial contribution to SBS98 remains a mechanistic hypothesis requiring future epidemiological validation.

### Decomposition of PanCancer genome data uncovers novel signature associations

To systematically map our experimental signatures onto human malignancies, we analyzed a pan-cancer dataset comprising 11,428 tumors from the PCAWG, TCGA, and Mutographs cohorts. We performed a decomposition of the NMF-extracted mutational profiles against a combined reference framework consisting of endogenous COSMIC signatures and our expanded eSS compendium. This compendium incorporates the other identified experimental signatures, eSS50 and eSS51, as the empirical representatives for SBS7c and SBS17b, respectively (**Suppl. Figure 11**). Experimental signatures resolved a defined fraction of the pan-cancer mutational landscape, detecting eSS-like mutational processes in 3,443 tumors (30.1%) across 38 distinct experimental signatures (**Figure 6al**; **Suppl. Figure 16**). Per-sample eSS burden was highest in skin (median 21.8 mutations per megabase) and lung (7.7 mutations per megabase) and fell below 1.5 mutations per megabase in all other tissues (**Suppl. Figure 17a**). The dominant processes matched the expected tissue biology: an eSS27-like (solar-radiation) process in 91% of skin tumors, an eSS18-like process in 75% of lung tumors, and an eSS33-like process in 59% of lung tumors. (**Figure 6b**; **Suppl. Figure 17b**). In esophageal cancers, eSS51 (5-FU) contributed 1.9 mutations per megabase in 18% of the samples. In head and neck cancers, eSS17 (B[a]P/BPDE), eSS18 (4NQO) and eSS33 (AAI/DBPDE) each contributed between 1.3 and 2.2 mutations per megabase (**Figure 6b**; **Suppl. Figure 17b**).

**Figure 6.**
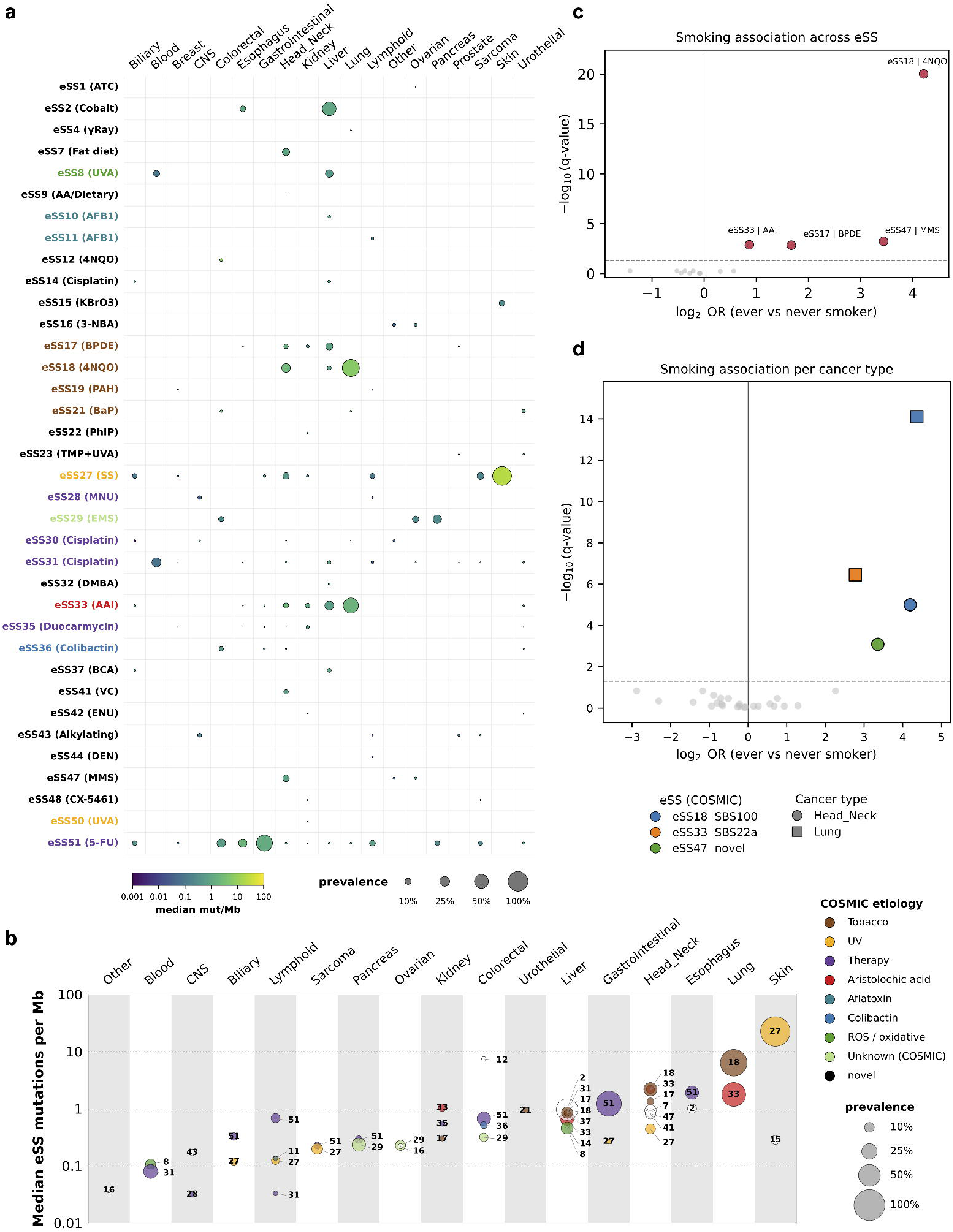
Pan-cancer attribution of the experimental SBS signatures. **(a)** Detection of the 49 eSS across 11,428 PCAWG, TCGA, and Mutographs tumors. Dot-matrix of each eSS (rows, labeled by dominant training mutagen) by cancer type (columns); dot size denotes carrier prevalence within the cancer type and dot color the median eSS burden (mutations/Mb). **(b)** Prevalence and burden of eSS by cancer type. Bubble plot restricted to eSS-cancer type pairs in which at least 2% of samples carry the eSS at ≥ 5% of total mutation burden. Bubble size denotes carrier prevalence and color the median carrier burden (mutations/Mb). **(c)** Volcano plot of the cancer type- and cohort-adjusted logistic association between eSS detection and smoking status. **(d)** Per-cancer type localization of the smoking signal. Within-tissue, cohort-adjusted associations confine the smoking enrichment to lung and to head and neck cancers.

Further, we identified four experimental signatures associated with smoking after adjusting for age, gender, cancer type, and cohort (**Figure 6c**), all enriched in ever-smokers within the smoking-annotated subset of 4,951 tumors (2,892 ever-smokers, 2,059 never-smokers). The tobacco signatures eSS17 (B[a]P; log_2_ OR 1.67, *q* = 1.4 x 10^-3^) and eSS18 (4NQO; log_2_ OR 4.21, *q* = 9.7 x 10^-21^) were recovered as expected in smokers and served as positive controls. The alkylating signature eSS47 (MMS and DMS; log_2_ OR 3.44, *q* = 5.7 x 10^-4^) was also enriched in smokers, reflecting a tobacco alkylating carcinogen that produces T>A mutations in A[T>A]N and T[T>A]T contexts, in addition to T[T>C]T mutations. The fourth signature, eSS33-like (AAI and DBPDE; log_2_ OR 0.87, *q* = 1.3 x 10^-3^), implicates DBP and DBPDE exposures rather than AAI, which accounts for 25% of the total signature attribution. Because DBP is a tobacco smoke PAH whose diol-epoxide metabolite induces an AAI-like T>A profile(*45*), this enrichment is likely driven by tobacco-derived PAH exposure (**Figure 6c**; **Suppl. Figure 17c**).

Per-cancer type logistic regression localized the smoking signal to lung and head and neck cancers (**Figure 6d**). eSS18 was enriched in ever-smokers in both lung (log_2_ OR 4.36, *q* = 8.0 x 10^-15^) and head and neck (log_2_ OR 4.19, *q* = 1.0 x 10^-5^). eSS33 was enriched in lung (log_2_ OR 2.78, *q* = 3.5 x 10^-7^), and eSS47 in head and neck (log_2_ OR 3.35, *q* = 8.1 x 10^-4^).

### Independent validation in human organoids by single-molecule sequencing

We reanalyzed an independent mutagenesis screen in which primary human organoids were exposed to carcinogens and sequenced by NanoSeq, a single-molecule duplex technology that resolves mutations at orthogonal error rates to the bulk sequencing(*46*). The dataset comprised 215 profiles across five tissues (stomach, colon, kidney, pancreas, liver), eight donors, and 19 compounds tested with and without S9 metabolic activation, alongside 66 controls. Each organoid pattern was cleaned with *SigRescueR* and matched to the 49-signature atlas(*25*).

Canonical carcinogens recovered their derivation-stage experimental signatures at high fidelity independent of the tissue (**Figure 7a-b**; **Suppl. Figure 18-19**). AAI reproduced eSS33 at cosine similarity of 0.98 across 16 samples, B[a]P reproduced eSS21 at 0.97 across 32 samples, aflatoxin B1 reproduced eSS10 at 0.95 across 14 samples, and the methylating agents MNNG and methylazoxymethanol reproduced eSS28 at 0.96 across 11 samples. The alkylating direct mutagen ENU reproduced eSS44 at 0.93, and potassium bromate reproduced its oxidative signature eSS15 at 0.97 (**Figure 7a**; **Suppl. Figure 19**).

**Figure 7.**
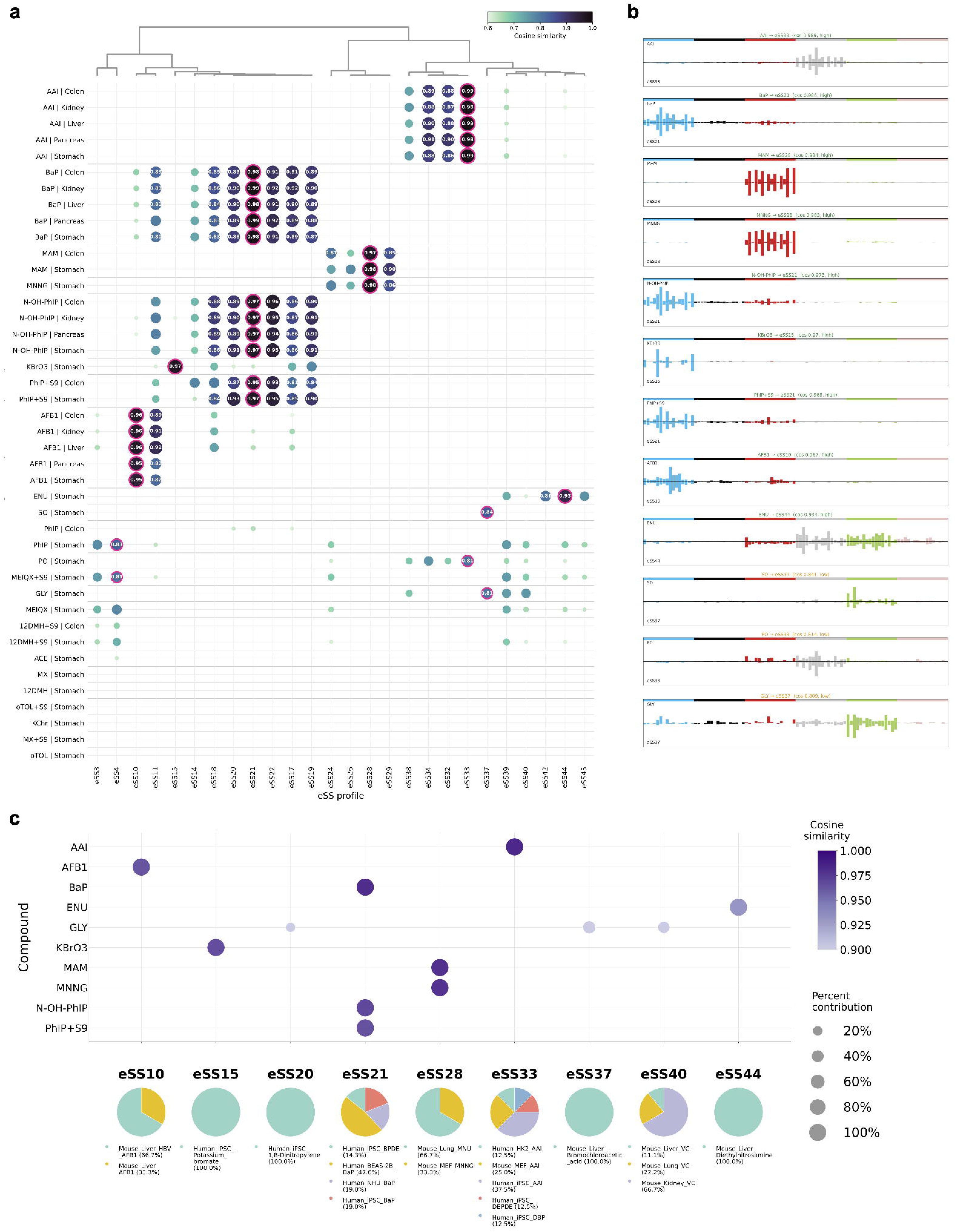
Independent validation using single-molecule NanoSeq 3D organoid data. **(a)** Clustered matrix of cosine similarity between each NanoSeq organoid mutational profile (rows, labeled by compound exposure and tissue) and the 49 eSS (columns). Dot color encodes cosine similarity (0.6 to 1.0). The best-matching eSS per exposure is outlined in magenta and labeled with its cosine value. **(b)** SBS96 plots comparing each organoid consensus profile (top) with its best-matching eSS (bottom). **(c)** Per-compound eSS decomposition dot plot. Dot color encodes cosine similarity (≥ 0.90) and dot size represent the percent contribution (≥ 5%). Pie charts below summarize the composition of each assigned eSS.

PhIP requires cytochrome-P450 activation to form its DNA-reactive metabolite. Without S9 mix, PhIP generated 249 substitutions and no coherent signature (**Figure 7a**; **Suppl. Figure 19**). With S9 mix, or when the pre-activated metabolite N-hydroxy-PhIP was applied directly, the organoids reproduced eSS21 at cosine similarity of 0.95 to 0.97 and burdens up to 9,226 substitutions, an increase of more than an order of magnitude. The PhIP signature in iPSC, eSS22, matched both activated forms of PhIP with a cosine similarity between 0.93 to 0.96 across tissues (**Figure 7a-b**; **Suppl. Figure 19**). Glycidamide, the reactive epoxide metabolite of the dietary contaminant acrylamide(*47*), was tested only in stomach organoids and it did not reproduce its compendium signature eSS39. The stomach profile matched eSS39 at only 0.750, below eSS37 (bromochloroacetic acid, 0.81). SigProfilerAssignment reconstructed it from eSS37 (45%) and eSS40 (36%) at cosine similarity of 0.87 (**Figure 7c**; **Suppl. Figure 20**).

Independent single-molecule sequencing validated these carcinogen signatures across exposure conditions, establishing that the mutational atlas generalizes across model systems and sequencing technologies, bounded primarily by exposure context.

## Discussion

Epidemiological models attribute 90% to 95% of cancer incidence to environmental and lifestyle factors(*48*). We have established the largest cross-species compendium of experimentally derived mutational signatures to date, capturing 49 distinct profiles induced by controlled environmental and chemical exposures. By pairing robust background attenuation via *SigRescueR* with stringent mutation-count thresholds and reproducible clustering, we effectively decoupled authentic exposure-driven signals from background noise(*25*). This resource significantly expands the experimental signature paradigm, which has predominantly relied on single-model frameworks, such as human iPSC surveys(*45*) or *C. elegans* DNA-repair mutants(*10, 49*), into a cross-species reference standard for interpreting mutational processes in human tumor cohorts.

A fundamental principle of this atlas is that a mutational signature represents the joint product of initial lesion chemistry and the host metabolic and repair pathways that process it(*10, 15, 50–52*). Genotoxic agents whose sequence specificity is dictated primarily by the initial chemical lesions, such as UV radiation and PAHs, consistently yield identical SBS profiles across diverse taxa, recapitulating their canonical COSMIC counterparts(*12*) (**Figure 3b-d**; **Figure 5a**; **Suppl. Figure 9**). Conversely, mutational processes dependent on specific repair machineries or complex xenobiotic activation reveal striking cross-model divergence. Host systems whose lesions are actively modified by mechanisms like MGMT, NER, or mismatch repair inherently reflect the repair capacity of the background organism and diverge accordingly. For instance, AAI failed to generate a reproducible signature in *C. elegans* (**Suppl. Figure 8**), a limitation stemming from restricted metabolic activation and the active clearance of bulky DNA adducts by specialized DNA-repair knockout backgrounds(*49*). Similarly, human models such as iPSCs frequently fail to recapitulate canonical signatures that demand complex xenobiotic metabolism and repair machineries(*45*). This divergence is particularly pronounced for therapeutic compounds like alkylating agents and platinum-based drugs that display high inter-species variance. These limitations underscore the critical necessity of utilizing human-matched tissue architectures for modeling these specific exposure classes.

The robustness of this paradigm is further validated by an independent duplex-sequencing screen of primary human organoids(*46*), which successfully reproduced the B[a]P, aflatoxin B1, AAI, and alkylating-agent signatures with cosine similarities ≥ 0.95 (**Figure 7**). Because this NanoSeq chemistry is completely independent of the clonal-expansion pipeline utilized in our cohort, this concordance supports platform-independent biological reproducibility rather than shared processing artifacts(*25*).

Decomposition of the 101 COSMICv3.6 signatures against our experimental anthology reinforced established etiologies while assigning putative experimental sources to previously uncharacterized signatures. In human cells, where NER clears aflatoxin N^7^-guanine and formamidopyrimidine adducts with comparatively lower efficiency than murine liver(*53*), the mutational spectrum shifts from SBS24 toward SBS29(*46, 54*). Consequently, a fraction of human SBS29 signatures aligns with an aflatoxin-driven process rather than tobacco chewing alone. However, in C>A-rich tissues like the lung, an SBS29 assignment most likely reflects spectral overlap with tobacco smoke or a mathematical fitting artifact, rather than genuine aflatoxin exposure (**Figure 5b**). Two previously unexplained orphan signatures were resolved into multi-agent processes rather than single-exposure events. SBS94 was robustly reconstructed from a majority of the B[a]P experimental signature (eSS17). B[a]P is a major constituent of tobacco smoke, yet SBS94 showed no association with smoking history(*55*). This points to a non-tobacco source of PAH exposure. Dietary intake of charred or burnt food is another plausible source, since it delivers B[a]P and related PAHs independent of smoking(*56*). This etiological link is also consistent with the isolated detection of SBS94 in colorectal cancers from Colombian cohorts(*55, 56*). Similarly, SBS95 was reconstructed from aflatoxin (eSS11) exposure, consistent with a dietary hepatotoxin exposure(*41*).

Our atlas also provides a putative chemical etiology for an additional cryptic signature. In liver malignancies, SBS101 was reconstructed from three mechanistically distinct genotoxins: DBP/DBPDE, bromochloroacetic acid, and glycidamide (eSS34, eSS37, and eSS39). DBP, a PAH present in tobacco smoke and high-temperature cooking, forms stable bulky adducts predominantly at adenine(*45, 57*). Glycidamide, the reactive epoxide of the high-temperature food contaminant acrylamide, forms N^7^-guanine and N^3^-adenine adducts that depurinate into mutagenic abasic sites (*47, 58, 59*). Meanwhile, bromochloroacetic acid, a drinking-water disinfection byproduct, generates a distinct hepatic mutational signature *in vivo* (*60, 61*), anchoring SBS101 in hepatic mutagenesis(*62*). Together, these reconstructions suggest a role for smoking and dietary mutagen exposures in liver carcinogenesis(*63, 64*). Next, the smoking signature SBS109 resolves into two distinct mutational chemistries (**Figure 5b**), including a bulky-adduct component (eSS12) and a methylating component (eSS47). This dual composition marks a specific alkylating tobacco exposure, distinct from the polycyclic-aromatic chemistry of the tobacco-smoke signature SBS4(*12, 65*). eSS47 detection was enriched in ever-smokers with head and neck cancer (*q* = 8.1 × 10^-4^), consistent with the SBS109 attribution(*65*) and supporting a role for a tobacco alkylating carcinogen, such as the tobacco-specific nitrosamines, in head and neck mutagenesis(*36, 66*).

In contrast, ionizing radiation, including gamma rays and heavy Fe-ions, produced no distinct SBS signature, instead resolving into organoid-derived signatures, eSS2 and eSS6. These matched the clock-like signatures SBS5 and SBS40, indicative of an accelerated cell proliferation and aging. Notably, rather than base substitutions, ionizing radiation imprints its primary genomic footprint through small deletions with junctional microhomology and balanced inversions(*10, 45, 67–70*).

Among the 24 reconstructed COSMIC signatures, 13 signatures (54.2%) displayed ambiguous non-unique linear combinations across alternative eSS (**Supplementary Table 10**). Importantly, this non-uniqueness reflected shared DNA damage mechanisms, identical lesion types, and common DNA damage response (DDR) pathways across exposures. This is particularly evident among signatures characterized by predominant C>A transversions and PAH exposure: SBS4, SBS24, SBS94, SBS95, and SBS100.

Collectively, the experimental signature attributions are mechanistic rather than compound specific, because agents sharing the same lesion type can converge on a single signature. Therefore, each eSS marks a class of DNA damage rather than a single chemical. The compendium samples 146 agents, a fraction of the human exposome, leaving chemical mixtures, chronic low-dose exposures, and endogenous processes largely not covered, so it functions as an evolving reference framework. This experimental catalog clarifies mutational processes operating in human cancers by linking environmental drivers to unannotated signatures, establishing a baseline for tracing tumor etiology and prioritizing carcinogens in public health prevention strategies.

## STAR Methods

### Data Curation

Publicly available datasets were obtained from the NCBI Sequence Read Archive (SRA) and European Genome-Phenome Archive (EGA) using the SRA Toolkit. Raw FASTQ files (5,349) were processed through an Ensembl variant calling pipeline. FASTQ files were aligned to the appropriate reference genome using BWA-MEM: mm10 for mouse data and GRCh38 for human data. Somatic mutations were identified from whole-genome sequencing (WGS) and whole-exome sequencing (WES) data using an ensemble approach across four independent variant callers: MuTect 2(*71*), VarScan 2(*72, 73*), Strelka 2(*74, 75*), and MuSE(*76*). Mutations supported by at least two of the four callers were retained(*24*). Variants were filtered to remove residual single nucleotide polymorphisms using dbSNP annotation via Ensembl Variant Effect Predictor (VEP). Additional filters excluded mutations shared across multiple samples and clustered events(*24*). The final set of high-confidence somatic mutations was carried forward for downstream analysis.

Pan-cancer WGS data made use of whole-genome sequencing (WGS) data generated by the Mutographs Cancer Grand Challenge project, an international, multi-cancer type resource designed to investigate geographic variation in mutational processes across populations with differing cancer incidence(*77*). Cancer type specific datasets and analyses from Mutographs used here include clear cell renal cell carcinoma (962 cases from 11 countries(*78*)), esophageal squamous cell carcinoma (552 cases from 8 countries(*79*)), head and neck cancer (265 cases from 8 countries(*65*)), and colorectal cancer (981 cases from 11 countries(*55*)).

Additionally, we incorporated WGS data from the Pan-Cancer Analysis of Whole Genomes (PCAWG) consortium(*80*), and The Cancer Genome Atlas (TCGA) Pan-Cancer Analysis Project(*81*).

Whole-genome sequencing data, mutational catalogues, and associated clinical/epidemiological metadata for the Mutographs cohorts are deposited in the European Genome-Phenome Archive (EGA) under study accessions EGAS00001003542 (clear cell renal cell carcinoma2), EGAS00001002725 (esophageal squamous cell carcinoma(*79*)), EGAS00001005450 (head and neck cancer(*82*)), and EGAS00001003774 (colorectal cancer(*55*)). PCAWG-generated alignments, somatic variant calls, annotations, and derived datasets(*80*) are available for general research use through the ICGC Data Portal at (http://dcc.icgc.org/pcawg/); most molecular, clinical, and specimen data are open-tier, while identifying data such as germline alleles require dbGaP (TCGA portion) or ICGC DACO (ICGC portion) approval. TCGA data are also available through the Genomic Data Commons (https://gdc.cancer.gov)(81).

### Data Processing

For human samples originally aligned to GRCh37, variant coordinates were converted to GRCh38 using LiftOver from Ensembl. Variant Call Format (VCF) files were pre-processed to ensure consistent formatting across datasets. For mouse WGS data, background signatures were detected and removed in two steps (**Suppl. Figure 3**). Non-negative matrix factorization (NMF) was applied to characterize the background(*41*). Background removal was then performed using *SigRescueR*(*25*), which subtracts baseline contributions per sample based on matched controls using default settings. Briefly, Bayesian inference was configured to run four Markov Chain Monte Carlo (MCMC) chains with 2,000 warm-up iterations followed by 4,000 sampling iterations per chain; the lower bound of the 95% credible interval (2.5th percentile) of the inferred exposure profile served as the representative estimate(*25*). The larger before-vs-after gap in human reflects higher endogenous and culture-associated background in human *in vitro* models. iPSC and immortalized lines undergo clonal expansion and extended passaging, so the multi-control *SigRescueR* framework subtracts a larger absolute count. Short-generation *C. elegans* and single-generation in vivo tumors carry less background and change less after cleanup.

In the *C. elegans* dataset, VCFs from all strains belonging to the same treatment group were aggregated to increase statistical power for downstream analyses. Mutational signature extraction and assignment were performed using the SigProfiler suite (SigProfilerMatrixGenerator, SigProfilerExtractor, SigProfilerAssignment), as described previously(*41, 83–85*).

### Hierarchical Clustering of Multi-Species Experimental Mutational Signatures

Mutational spectra data were obtained from five species: human, mouse, rat, chicken, and *Caenorhabditis elegans* (*C. elegans*). To test the stability of individual mutational profiles, we subjected the harmonized SBS96 dataset to Poisson resampling (**Suppl. Figure 6a**). This simulation modeled random sampling variability and unsystematic noise, revealing that statistical confidence in a given profile scales directly with total mutational burden. Profiles were then filtered to include only samples with ≥307 total mutation counts, a threshold established by Poisson resampling analysis (**Suppl. Figure 6a**). Sample names were prefixed with species identifiers, and all datasets were combined into a single matrix with profiles normalized to sum to 1.0.

Hierarchical clustering used cosine distance (1 − cosine similarity) with agglomerative clustering and average linkage (scipy.cluster.hierarchy v1.13.1). A cosine distance threshold of 0.1 (90% cosine similarity) defined discrete clusters. A two-stage clustering approach was applied, with secondary clustering of clusters containing both aristolochic acid I (AAI) and dibenzo[a,l]pyrene diol-epoxide (DBPDE) signatures to achieve optimal separation of the distinct mutagens. Clusters were retained if they contained ≥3 eSS samples.

Consensus signature profiles were calculated using context-weighted averaging: for each mutational context, sample contributions were weighted proportionally to raw mutation counts for that context, then applied to normalized signature values. Consensus profiles were renormalized to sum to 1.0 and compared against COSMICv3.6 using cosine similarity. Profiles with similarity ≥0.85 to any COSMIC signature were classified as similar to known signatures; profiles with similarity <0.85 were classified as potentially novel patterns. Consensus signatures were visualized as SBS-96 plots using the SigProfilerPlotting tool (v1.4.1).

Pairwise cosine similarity among the 49 eSS was computed on the 96-channel SBS profiles, and an undirected graph was built with an edge at cosine similarity ≥ 0.90. Connected components of at least two members defined 10 reproducible clusters, rendered by Fruchterman-Reingold force-directed layout.

### COSMICv3.6 decomposition against the experimental signatures

COSMICv3.6 SBS signatures were decomposed against the 49 eSS with SigProfilerAssignment using the eSS profiles as a custom signature database. For each COSMIC signature the algorithm added eSS components layer by layer and retained the composition maximizing cosine similarity to the target, reporting the final L2 error and cosine. Reconstructions with cosine >=0.90 were called explained, and eSS contributing less than 5% were dropped to suppress noise. Per-mutation-class cosine similarity between each eSS and COSMICv3.6 was computed by restricting both profiles to the 16 channels of each of the six substitution classes.

### Robustness and identifiability of the COSMIC reconstructions

SigProfilerAssignment reconstructed 48 of 101 COSMIC signatures at cosine similarity ≥ 0.80 and declared the remaining 53 novel relative to the eSS panel. For each of the 48 reconstructed signatures, all 18,424 three-eSS combinations were refit using SigProfilerAssignment, retaining combinations in which all three eSS carried ≥ 5% weight. Combinations whose reconstruction cosine reached the SigProfilerAssignment cosine were counted as near-degenerate alternatives. A reconstruction was classified robust when it admitted ≤ 3 such alternatives and its selected eSS were not collinear (maximum pairwise cosine < 0.85), and ambiguous otherwise. Component stability was tested by drawing 1,000 multinomial resamples of each 96-channel profile at a depth of 10,000 mutations (seed 42), refitting each resample against the 49 eSS, and recording the selection frequency of each eSS at > 5% weight.

### Pan-cancer eSS attribution and association analysis

The eSS reference was attributed to *de novo* signatures extracted using SigProfilerExtractor tool from 11,428 pan-cancer whole-genome tumors across PCAWG, TCGA, and Mutographs cohorts with SigProfilerAssignment, Per-sample activities were converted to relative contributions. Smoking association was tested for each eSS between ever-smokers and never-smokers within the 4,951 smoking-annotated tumors (2,892 ever-smokers, 2,059 never-smokers) with a two-sided Mann-Whitney U test. Logistic regression adjusting for age, gender, cancer type, and cohort was performed as follows:

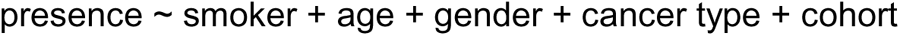

Benjamini-Hochberg FDR was computed across the eSS. Calls required q < 0.05. *De novo* signatures were extracted per cancer type and decomposed against the hybrid eSS reference with SigProfilerAssignment, applying increased penalties for reconstruction: nnls_add_penalty = 0.10, nnls_remove_penalty = 0.01, and initial_remove_penalty = 0.05.

### Independent organoid validation by single-molecule (NanoSeq) sequencing

An independent carcinogen mutagenesis screen in primary human organoids was reanalyzed to test signature reproducibility outside the derivation models(*46*). The dataset comprised 215 SBS96 profiles across five tissues (stomach, colon, kidney, pancreas, and liver), eight donor lines, and 19 compounds tested with and without S9 metabolic activation, alongside 66 controls. Background signal was removed with *SigRescueR*, yielding 149 treatment-attributable per-sample profiles(*25*). For each exposure, a mutation-burden-weighted average SBS96 profile was computed across replicates, weighting each sample by its mutation count so that low-count profiles did not dominate the aggregate. Compound-level profiles were pooled across tissues; pooling was justified by a homogeneity check in which every multi-tissue compound with signal reached cross-tissue cosine ≥ 0.95. Condition-level profiles (compound by tissue) were retained where tissue resolution mattered. Each aggregated organoid profile was matched to the 49 eSS by cosine similarity. Cleaned profiles reproduced the source study per-compound signatures at cosine 0.91 to 1.00, confirming that background removal recovered the published profiles rather than distorting them. Decomposition of the organoid data, at both the individual sample and aggregated levels, against the 49 eSS signatures was performed using default settings in SigProfilerAssignment(*86*).

### Statistical Analysis

Tumor mutational burden (TMB) was calculated as somatic mutations per megabase: Human genome 3200 Mb / exome 30 Mb; Mouse genome 2800 Mb / exome 30 Mb; Rat genome 2750 Mb; *C. elegans* genome 100 Mb; Chicken genome 1200 Mb. Cosine similarity was used to assess similarity between extracted and reference signatures. Odds ratios with 95% confidence intervals quantified relative enrichment of mutation classes or signatures across treatment conditions. Analyses and visualizations were conducted in R (v4.4) and Python3 (pandas, numpy, scipy).

## Code and Data Availability

All code used to generate the figures and analyses reported in this manuscript is openly available on GitHub at: https://github.com/ZhivaguiLab/eSignature_MS/. Detailed instructions for reproducing the results are provided in the repository’s README file.

## Author Contributions

M.Z. and L.B.A. conceptualized the study. M.Z., S.S., and M.B. amassed data from biorepositories and publications and ran genomic analysis. M.Z., J.N.A., S.S., P.N., S.A., and Z.J. conducted data analyses. M.Z. and L.B.A. wrote the manuscript with input from all co-authors. All authors reviewed and approved the final version of the manuscript.

---

**Supplementary Figure 1.**
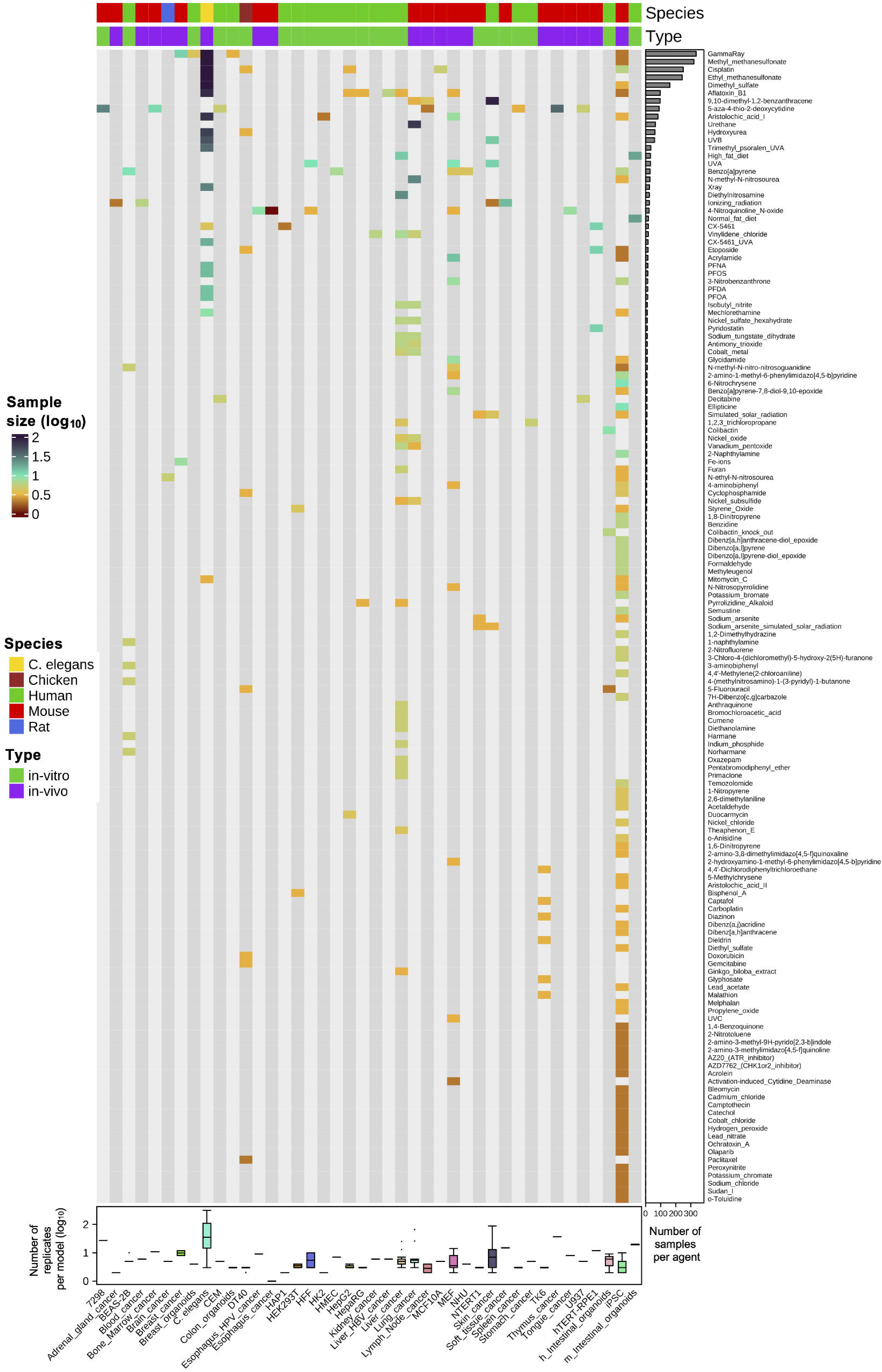
Heatmap of mutagen exposure data across experimental conditions. Columns represent individual experimental model systems, annotated by species and experiment type. Rows depict mutagenic agents, grouped by total sample count across models.

**Supplementary Figure 2.**
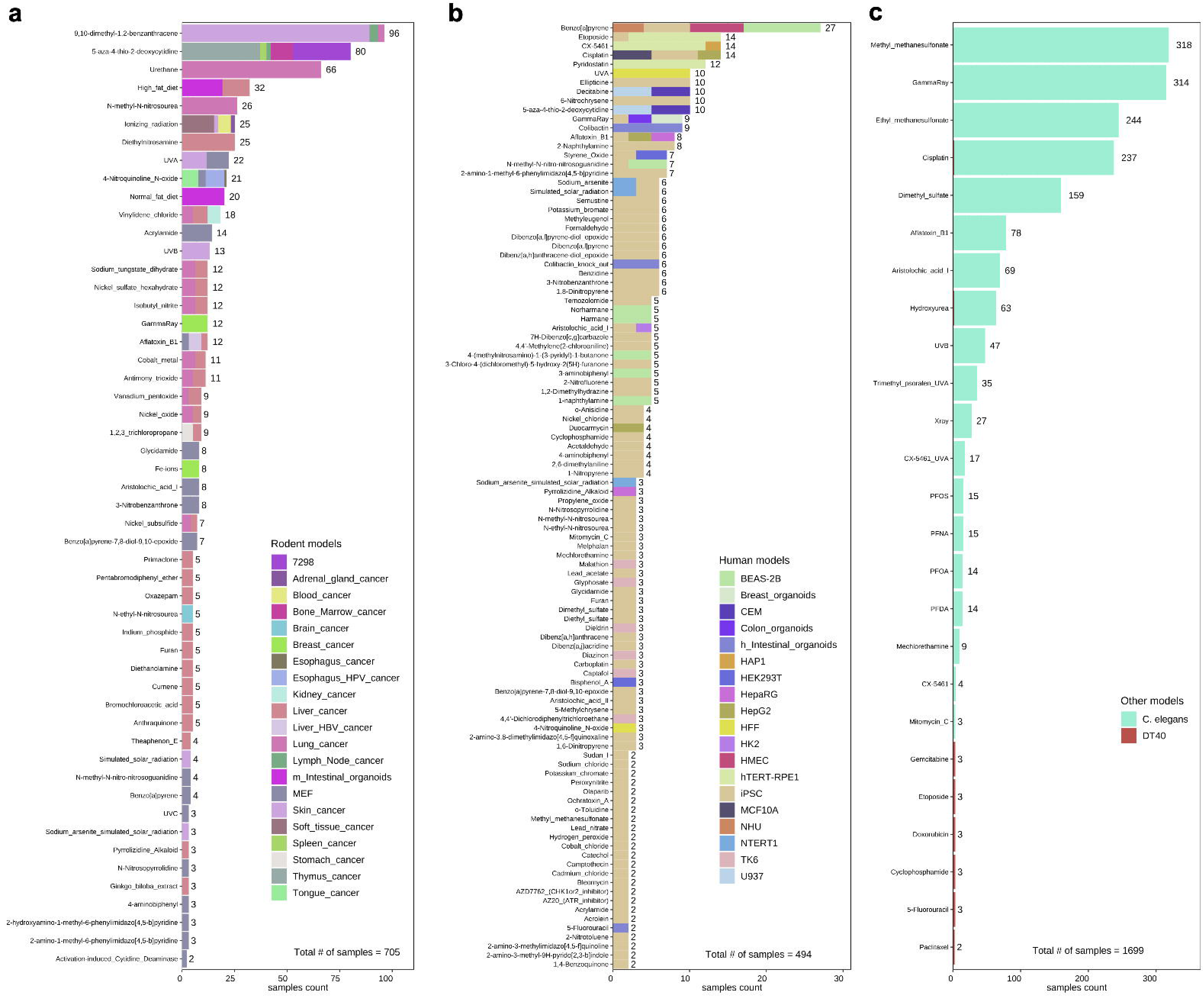
Distinct mutagenic agents tested across model systems and species. Panels show (a) rodents, (b) humans, and (c) other species, including *C. elegans* and chicken cell lines. Numbers represent the quantity of replicates per model system.

**Supplementary Figure 3.**
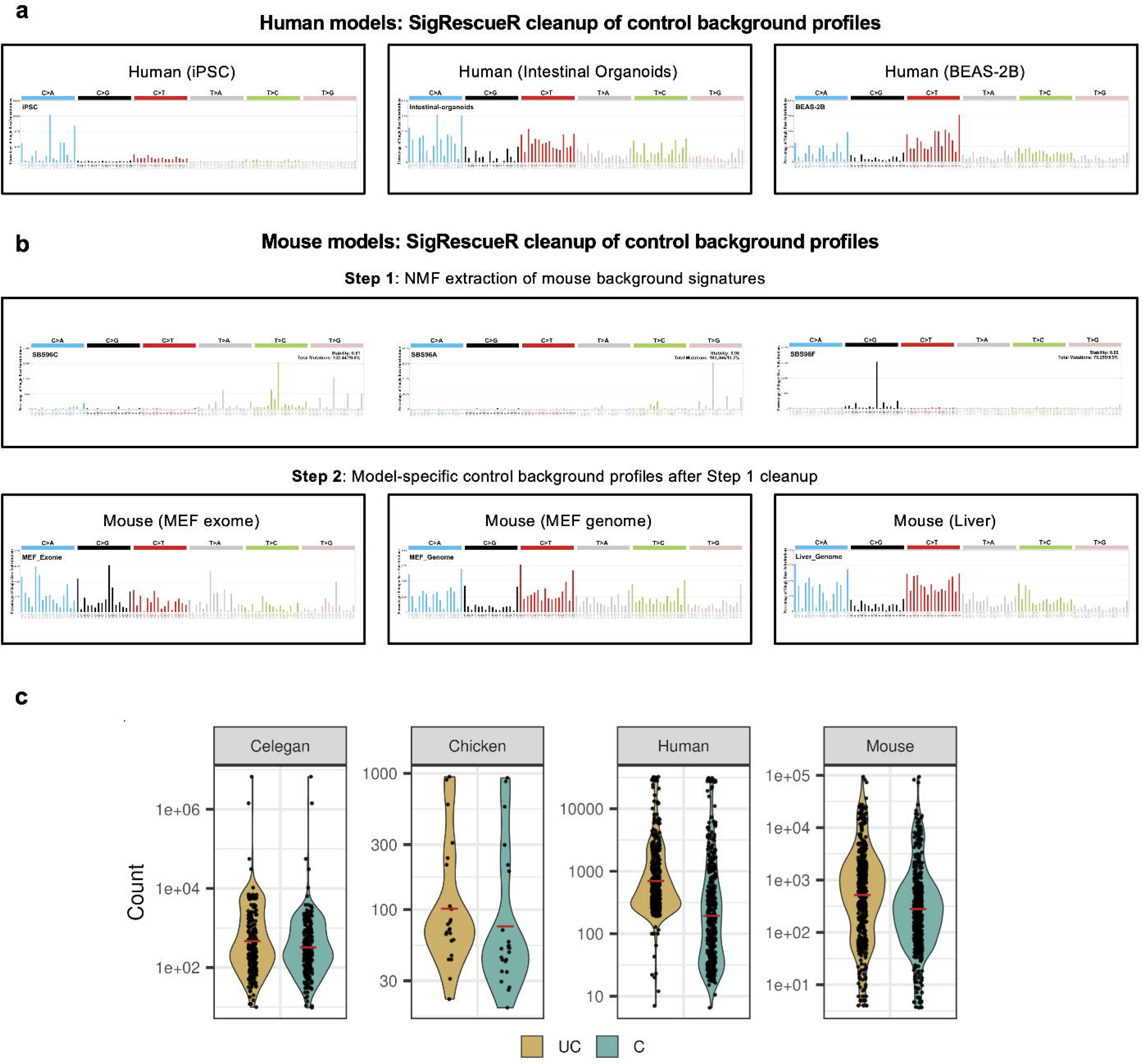
SigRescueR background filtering process in human cells and mouse models. Filtering workflow for (a) human cells and (b) mouse models. Mouse samples underwent a two-step cleanup strategy: in step 1, ubiquitous mouse-specific background processes (such as culture artifacts) were removed from exposed and control samples; in step 2, the remaining model-specific background profiles were subtracted using processed controls. (c) Number of SBS mutations before and after cleanup across species.

**Supplementary Figure 4.**
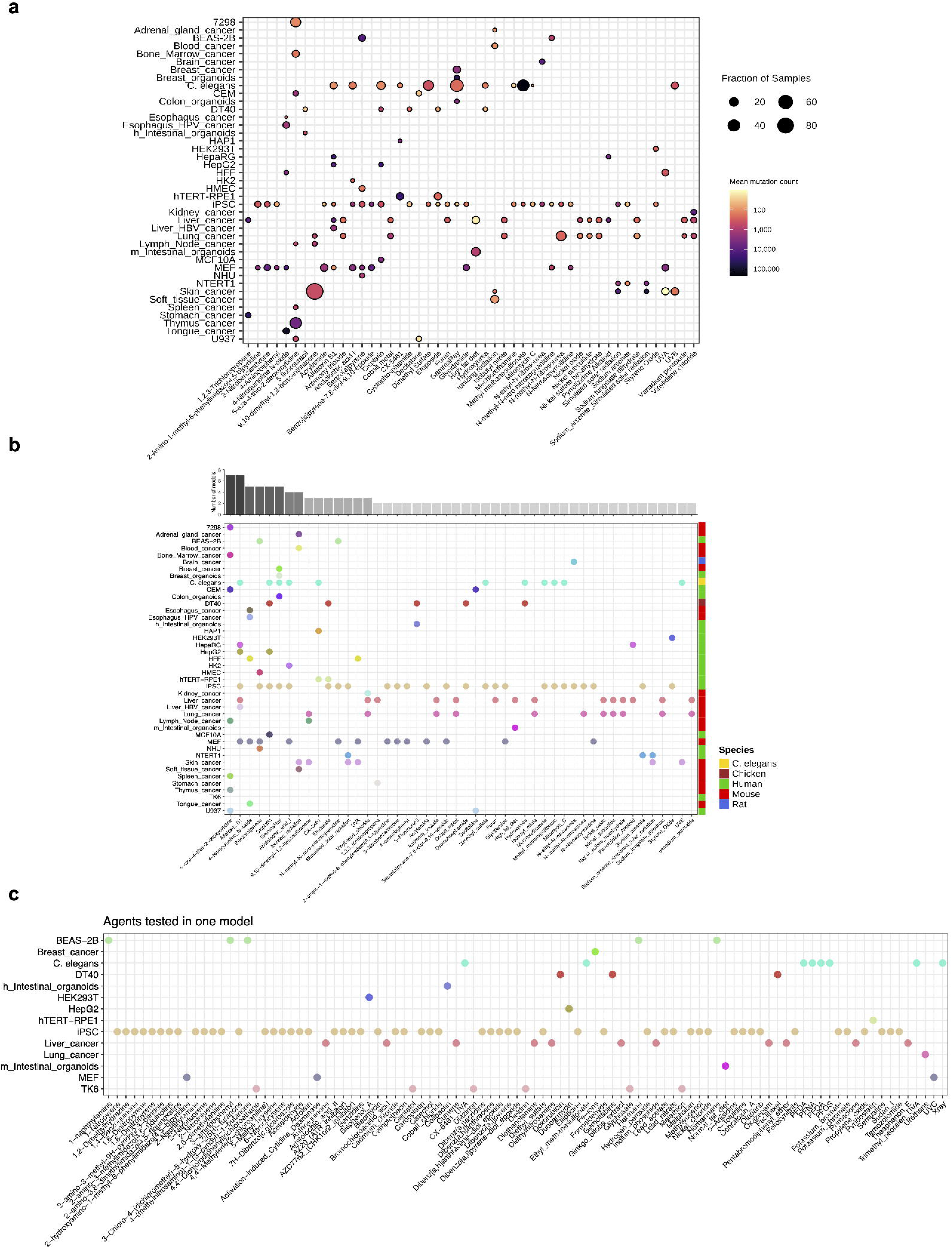
(a) Mutational burden per model system for agents tested in at least two model systems, with 48 compounds profiled across multiple species ranked by median burden. (b) Distribution of the 48 compounds shown per model system and compound. (c) Distribution of the remaining agents tested in only a single model system.

**Supplementary Figure 5.**
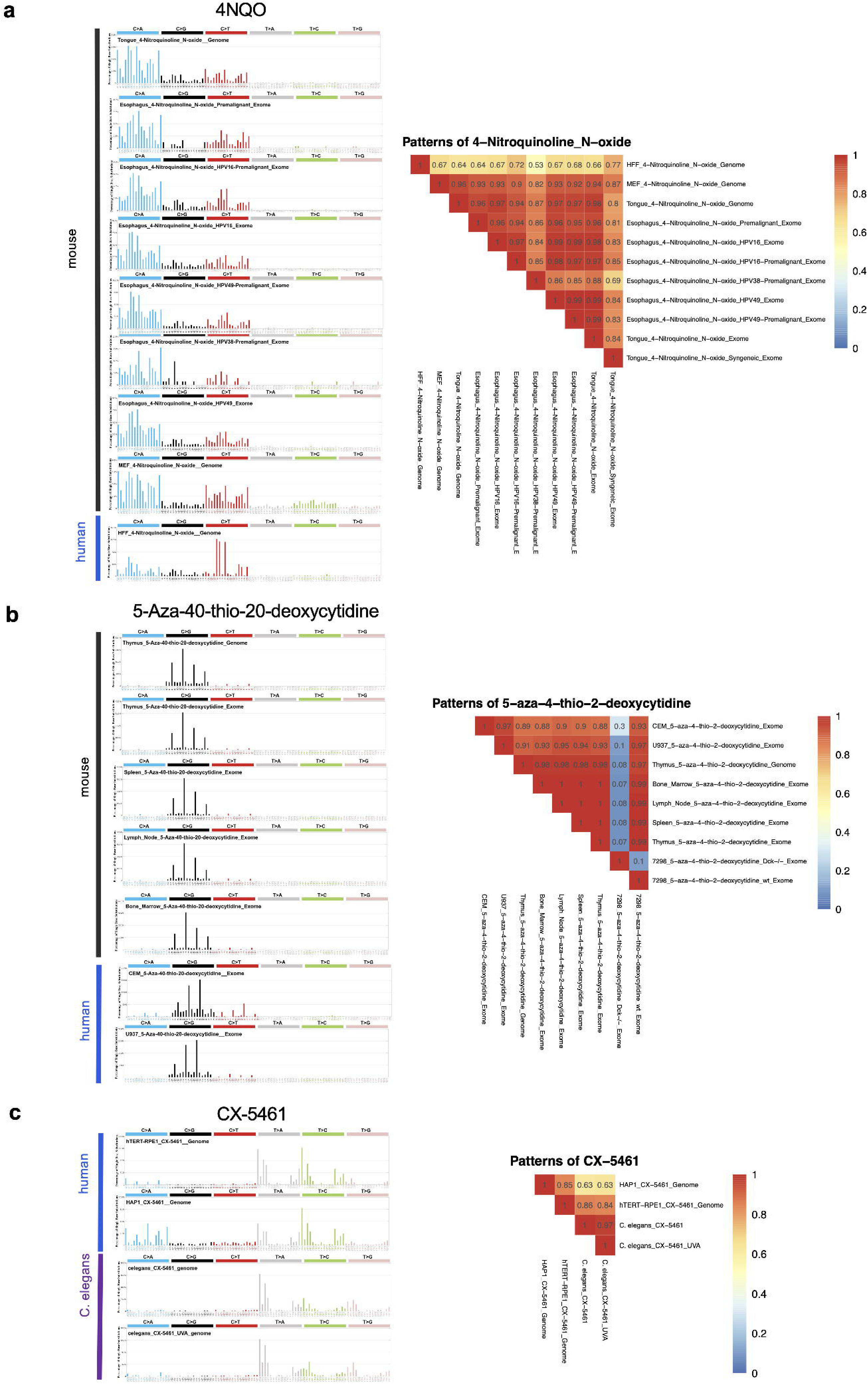
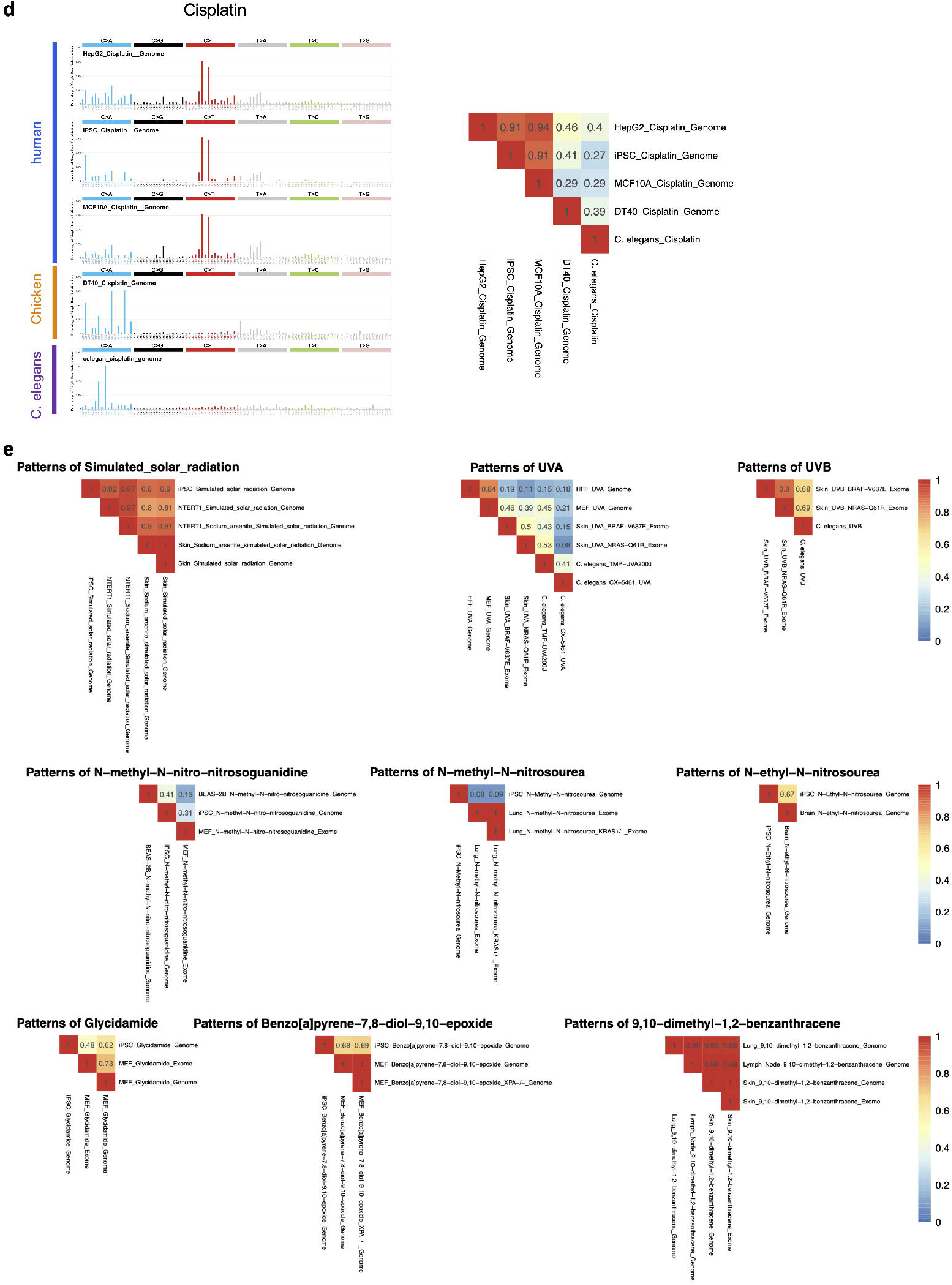
Reproducibility of SBS96 mutational signatures across model systems. SBS96 spectra and pairwise cosine similarities across model systems and species for (a) 4NQO, (b) ATC, (c) CX-5461, and (d) cisplatin. (e) Pairwise cosine similarities for additional exposures, such as solar radiation, UVA, and UVB. Shown are compounds tested in more than three model systems, with per-agent cross-model cosine similarity heatmap.

**Supplementary Figure 6.**
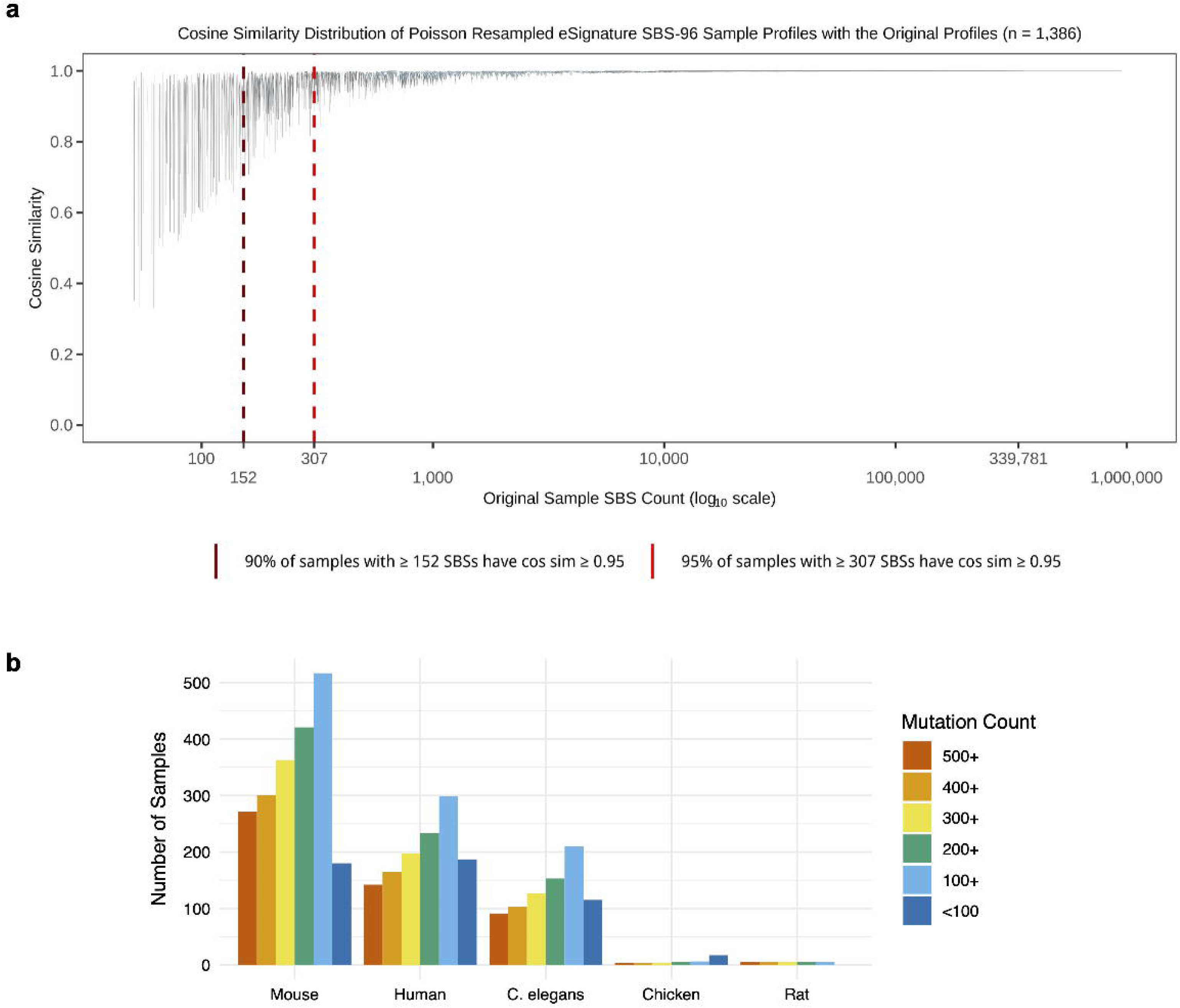
Robustness of the experimental SBS96 mutational profiles evaluated via Poisson resampling. (a) Violin plots illustrate the distribution of cosine similarities between 1,000 Poisson-bootstrapped replicates per sample and their corresponding original harmonized profiles (n = 4,282). The x-axis shows raw mutation counts for the original samples on a log_10_ scale. The dark red dashed line denotes the lowest mutation burden at which 90% of resampled iterations retain a cosine similarity ≥ 0.95 relative to the baseline profile. The bright red line marks the burden threshold required for 95% of iterations to achieve this level of concordance. (b) Fraction of profiles exceeding the threshold.

**Supplementary Figure 7.**
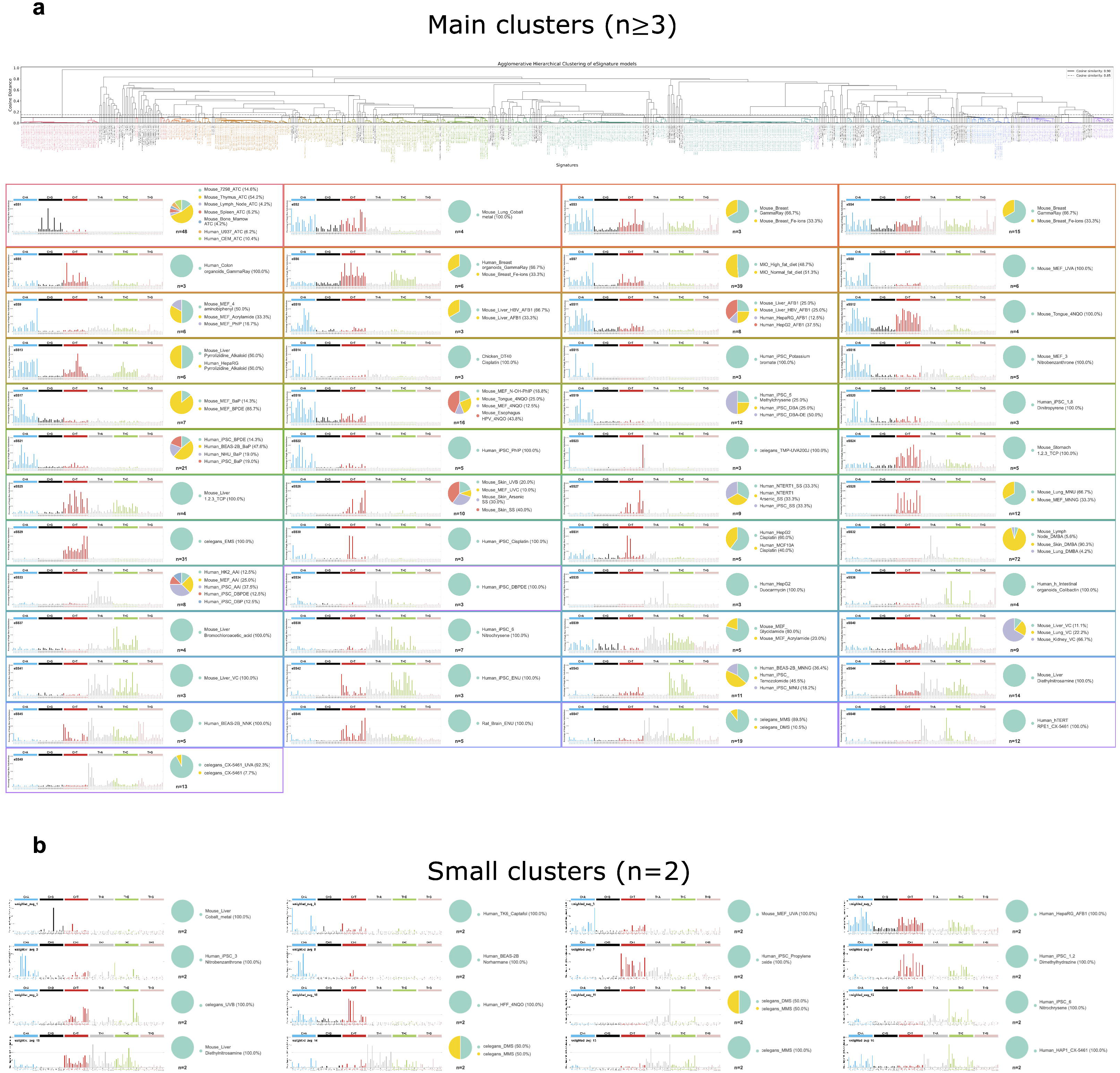
Hierarchical clustering of high-confidence profiles with cosine-distance dendrogram of the 671 high-confidence profiles defining the 49 eSS (≥ 3 samples per cluster). (b) Small clusters of n = 2 that did not meet the eSS reproducibility criterion.

**Supplementary Figure 8.**
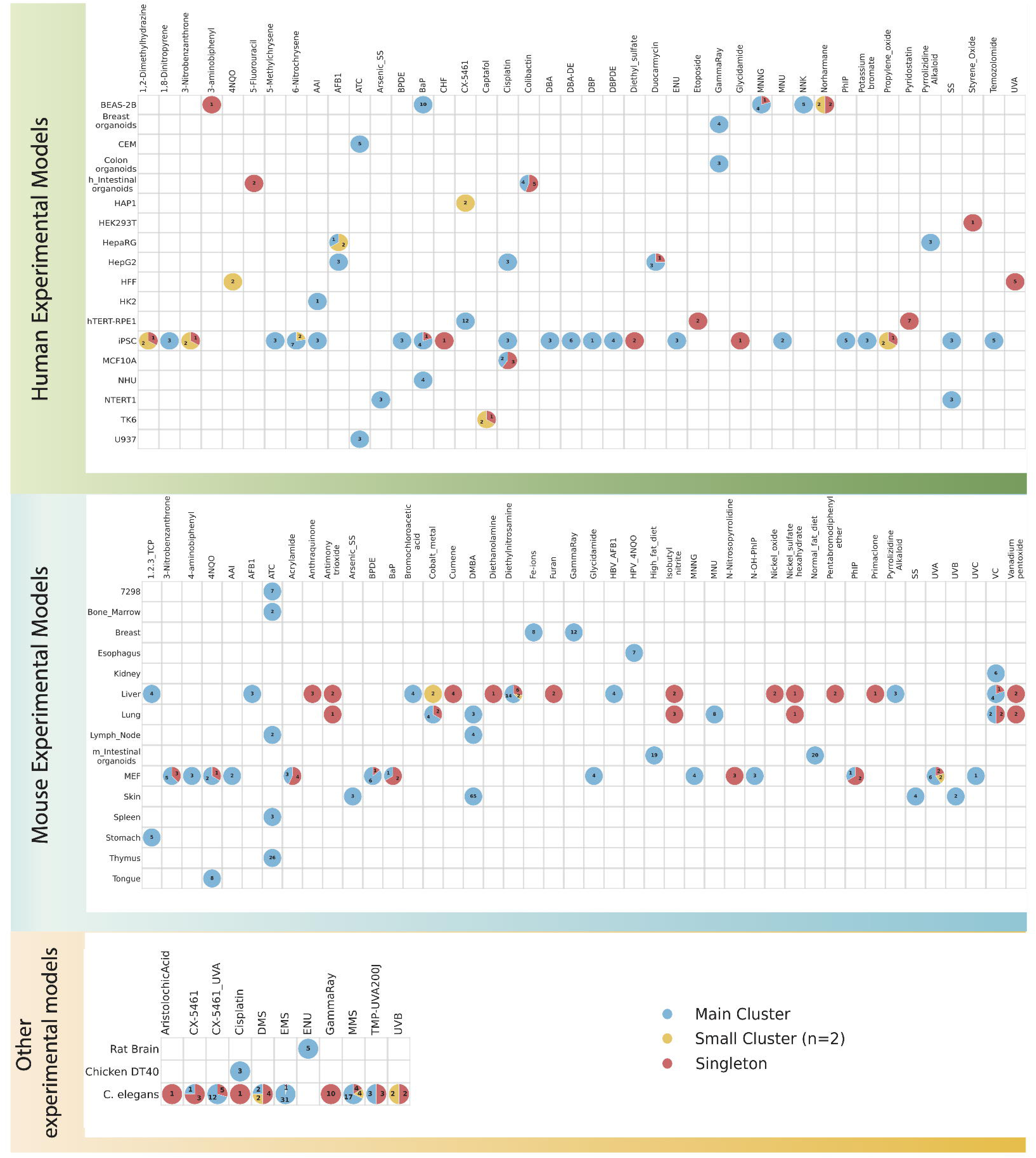
Sample distribution across cluster categories per model system. Heatmap overlaid with pie charts representing the number of samples per compound categorized as a main cluster (n ≥ 3), small cluster (n = 2), or singleton (n = 1) across model systems. Shown are human models (top panel), mouse models (middle panel), and other models (*C. elegans*, chicken cells, and rat tumors; bottom panel).

**Supplementary Figure 9.**
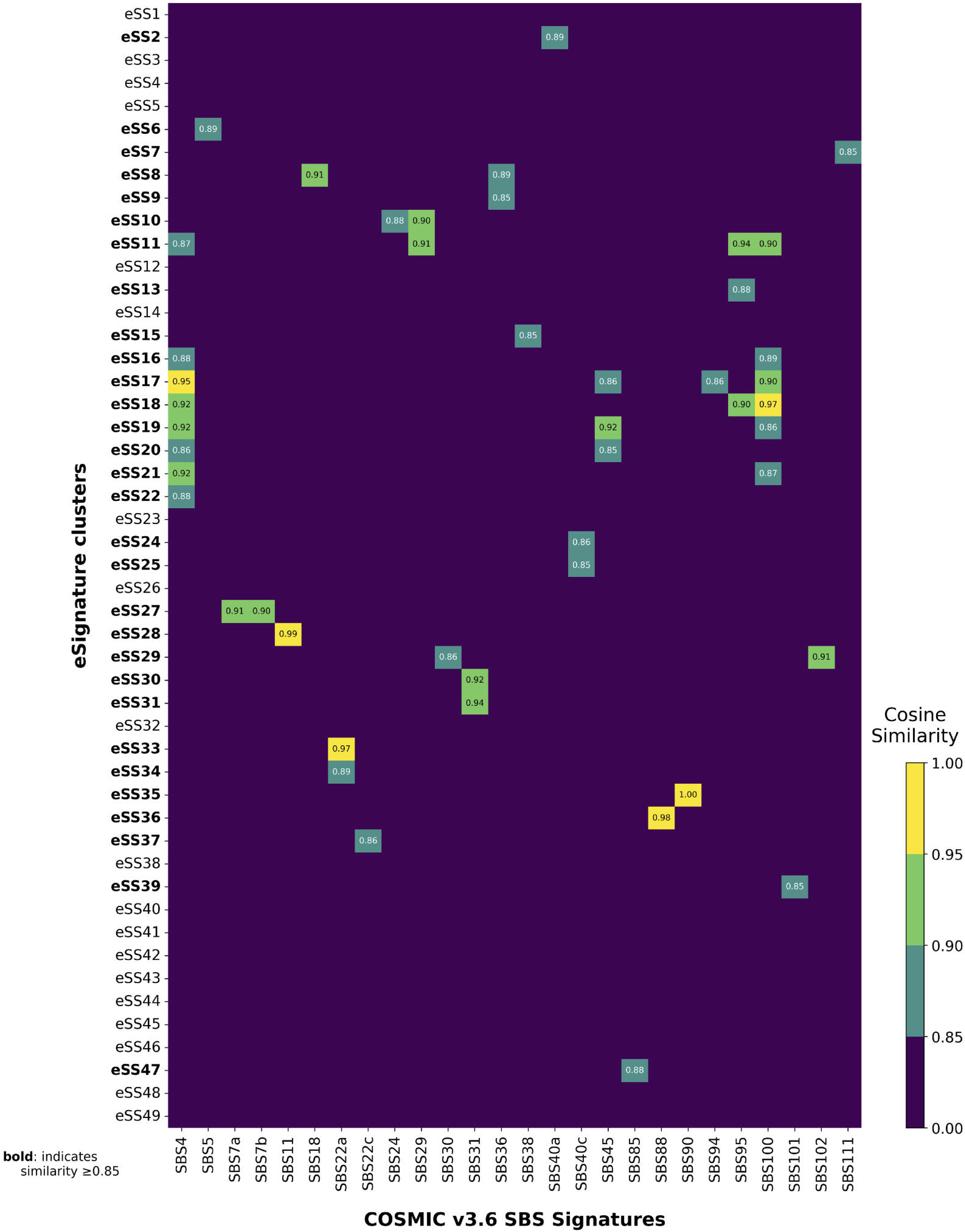
Heatmap of the cosine similarity between the 49 eSS and COSMIC v3.6, ranging from 0.85 to 1.

**Supplementary Figure 10.**
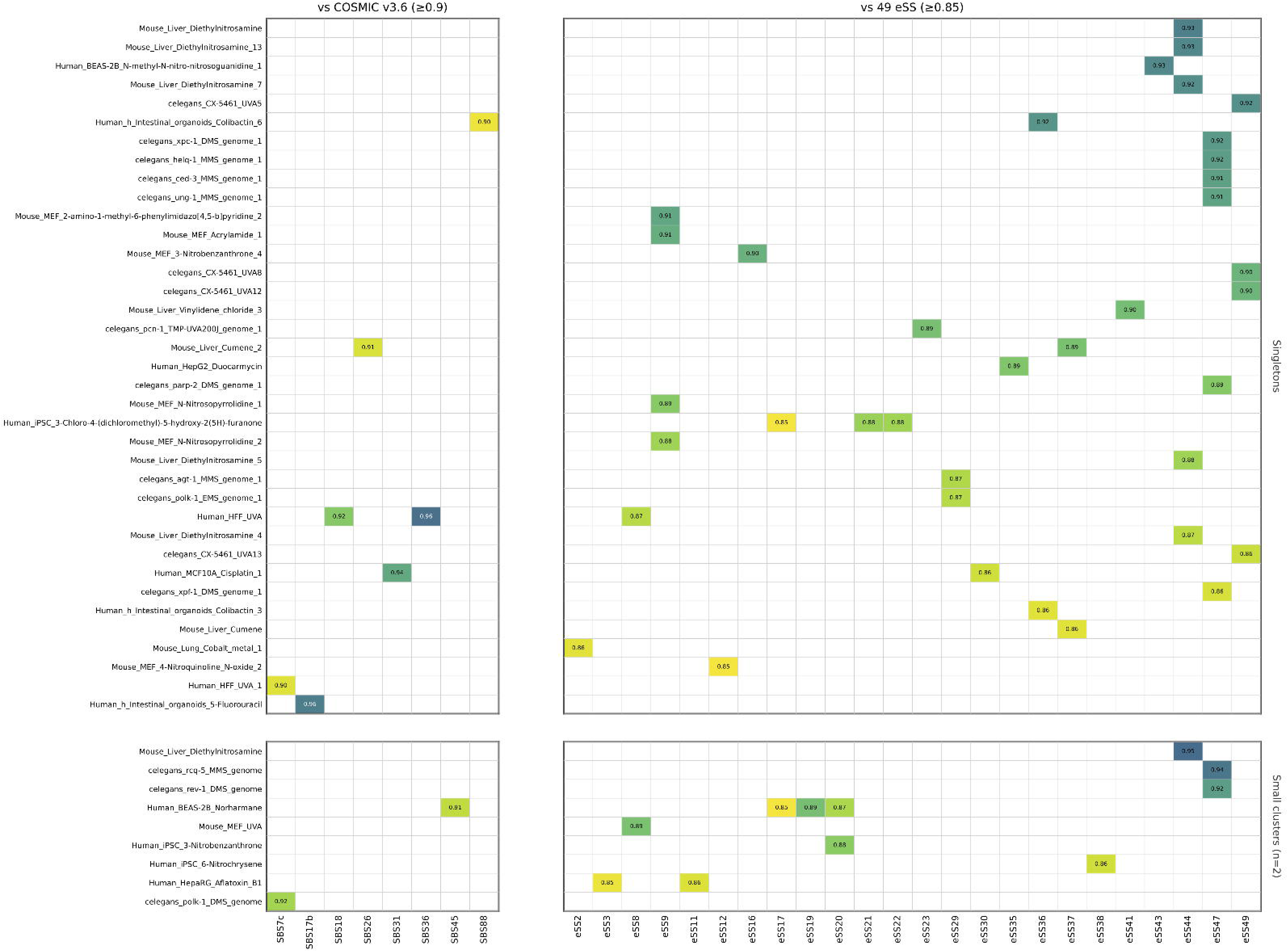
Cosine similarity of singleton and n = 2 profiles against both (a) the full COSMICv3.6 SBS database and (b) the 49 eSS, scrutinizing signal missed by the eSS anthology.

**Supplementary Figure 11.**
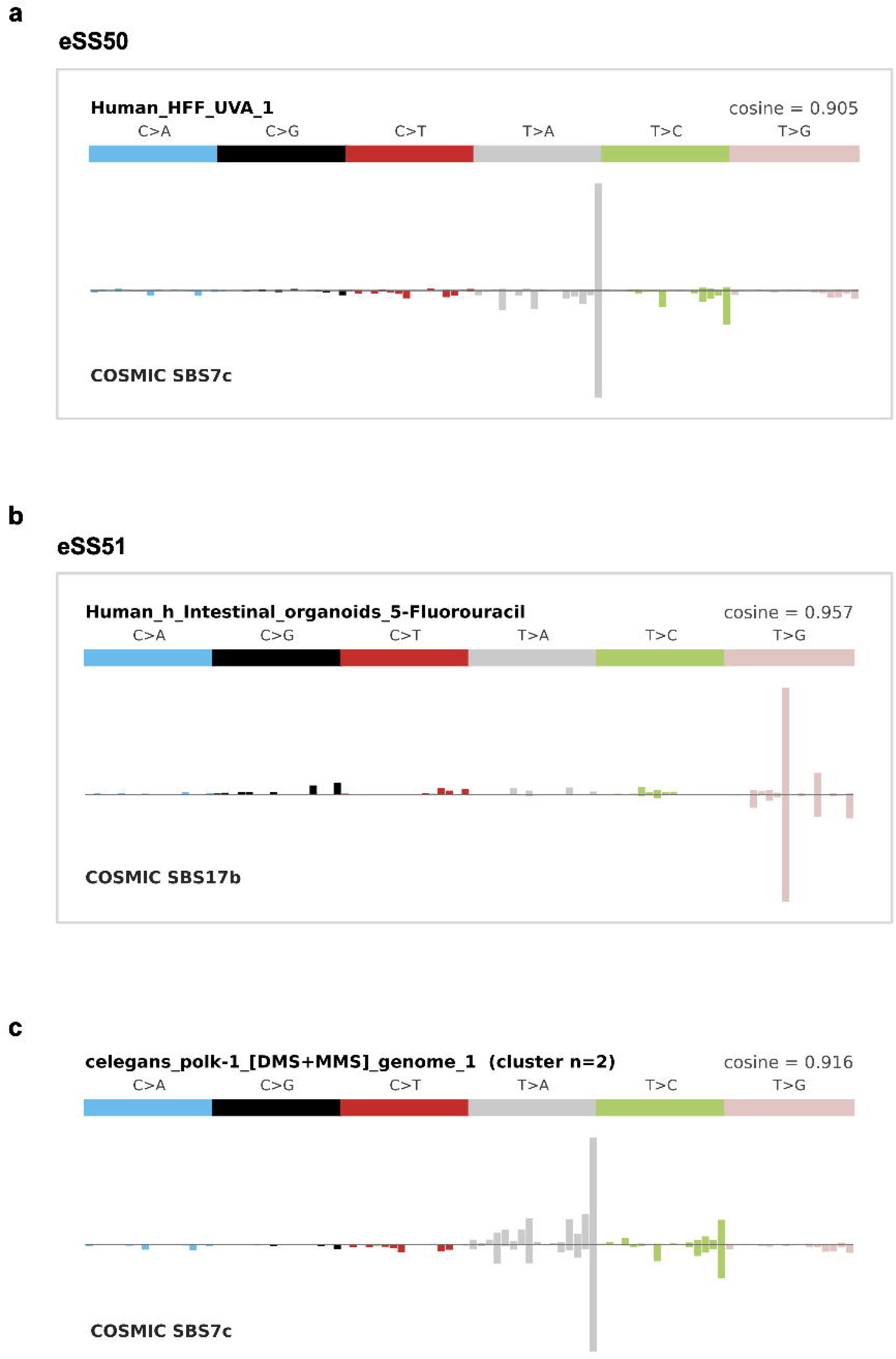
The three profiles reaching cosine ≥ 0.9 to a COSMICv3.6 signature while remaining outside the 49-eSS compendium: (a) UVA in HFF matching SBS7c, (b) 5-FU matching SBS17b, and (c) DMS and MMS in *C. elegans* matching SBS7c.

**Supplementary Figure 12.**
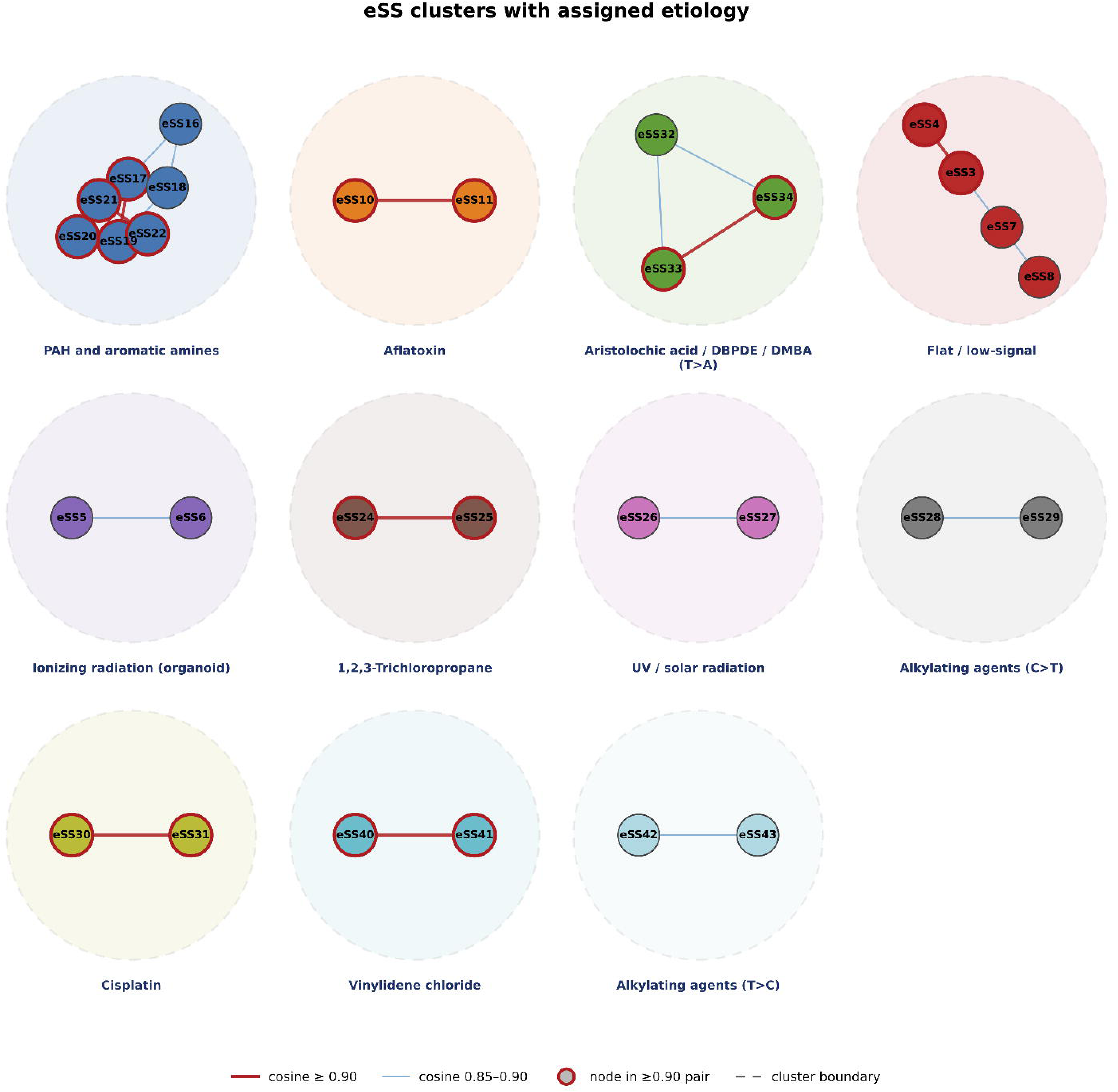
Pattern families of the 49 eSS. Network and cluster structure of the seven pattern families. The aflatoxin pair separates from the PAH and nitroaromatic core at cosine ≥ 0.90.

**Supplementary Figure 13.**
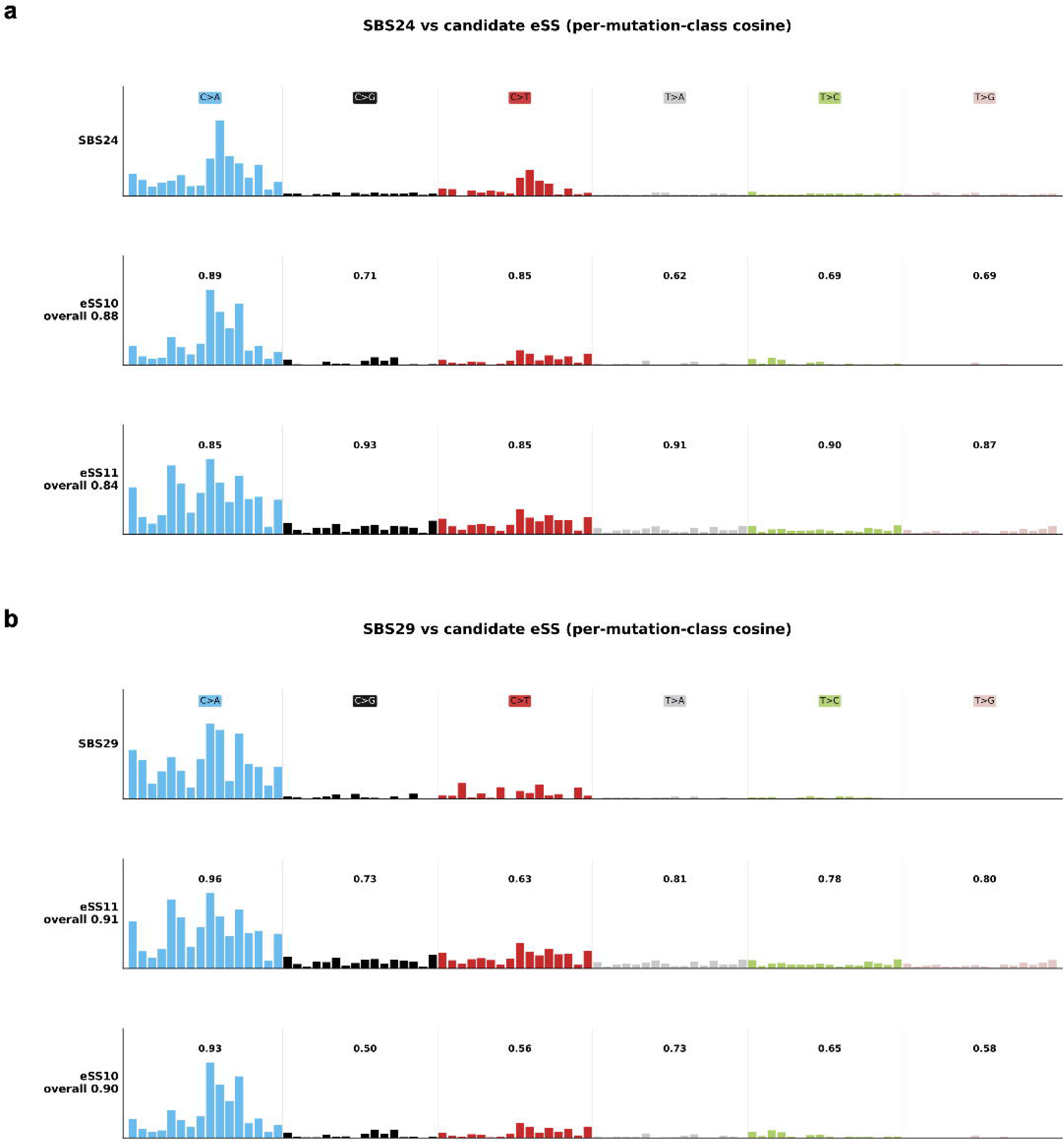
Decomposition of COSMIC signatures using aflatoxin eSS profiles. (a) Reconstruction of SBS24 from the mouse-liver aflatoxin signature eSS10. (b) Reconstruction of SBS29 from the human aflatoxin signature eSS11. Both panels display the per-mutation-class cosine similarity.

**Supplementary Figure 14.**
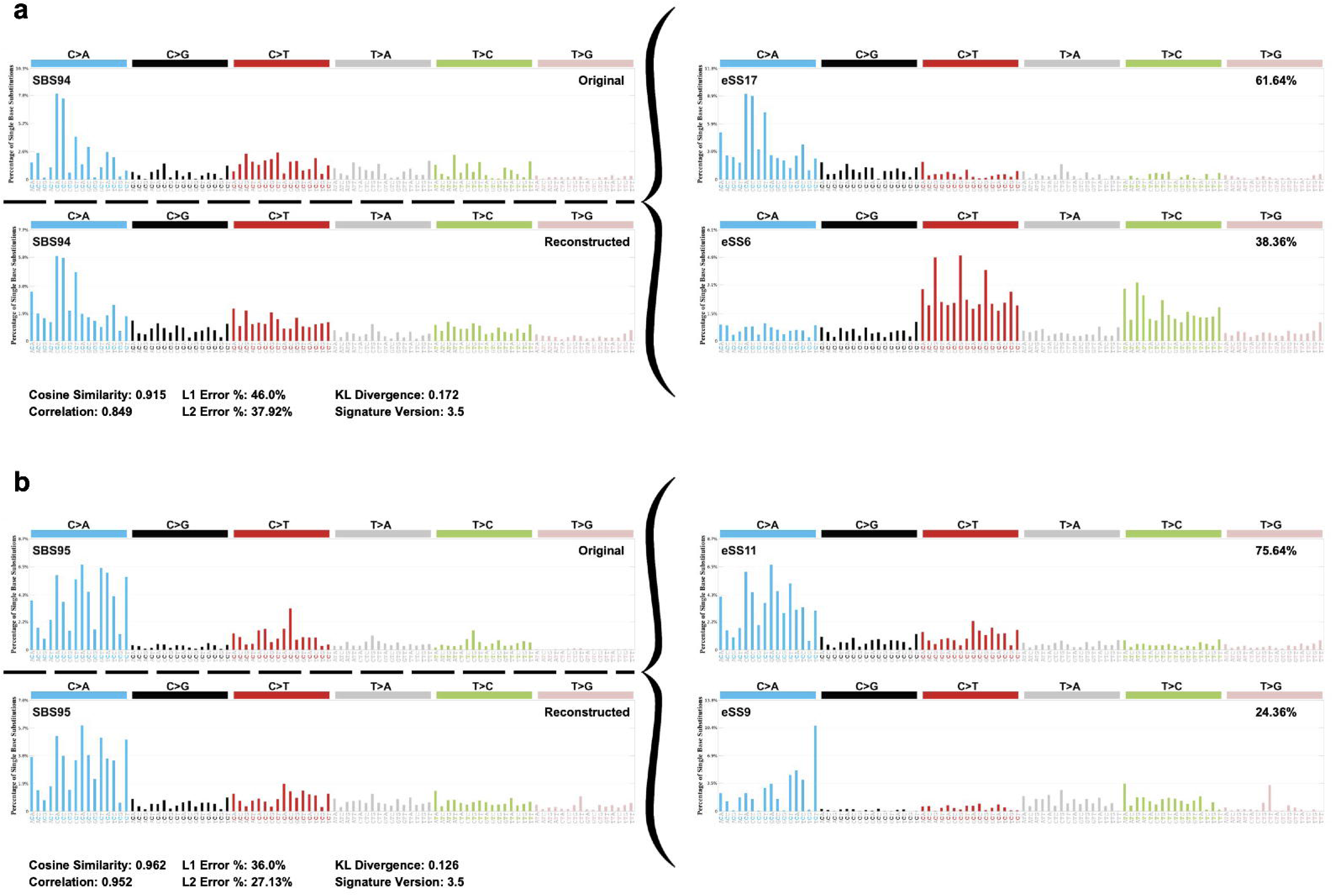
Decomposition of orphan COSMIC signatures. (a) SBS94 reconstructed from eSS6 and eSS17 (dietary PAH). (b) SBS95 reconstructed from eSS11 (aflatoxin and related mycotoxin).

**Supplementary Figure 15.**
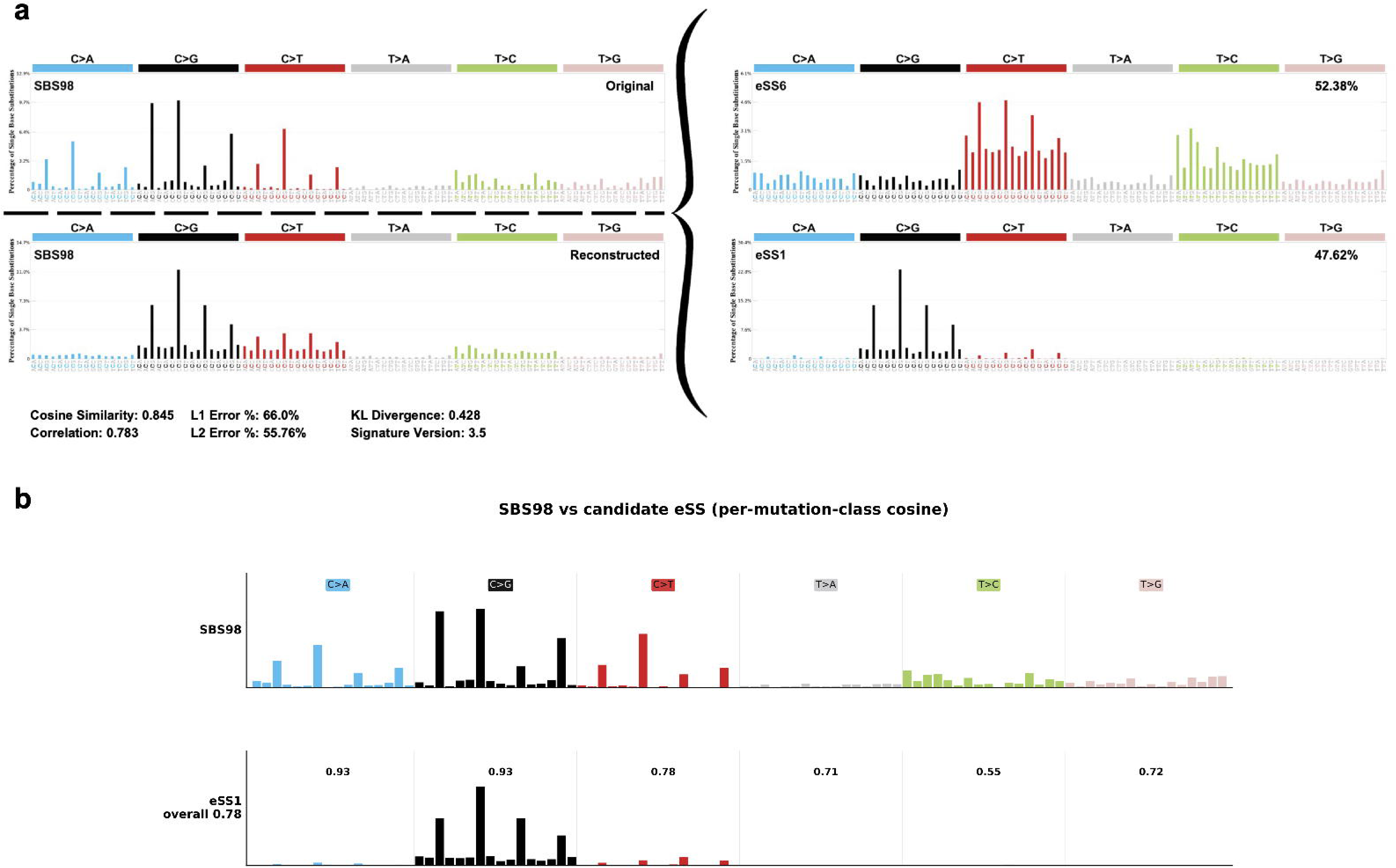
eSS1 and COSMIC SBS98. (a) NNLS decomposition of SBS98 against the eSS atlas. (b) Per-mutation-class cosine similarity between SBS98 and eSS1.

**Supplementary Figure 16.**
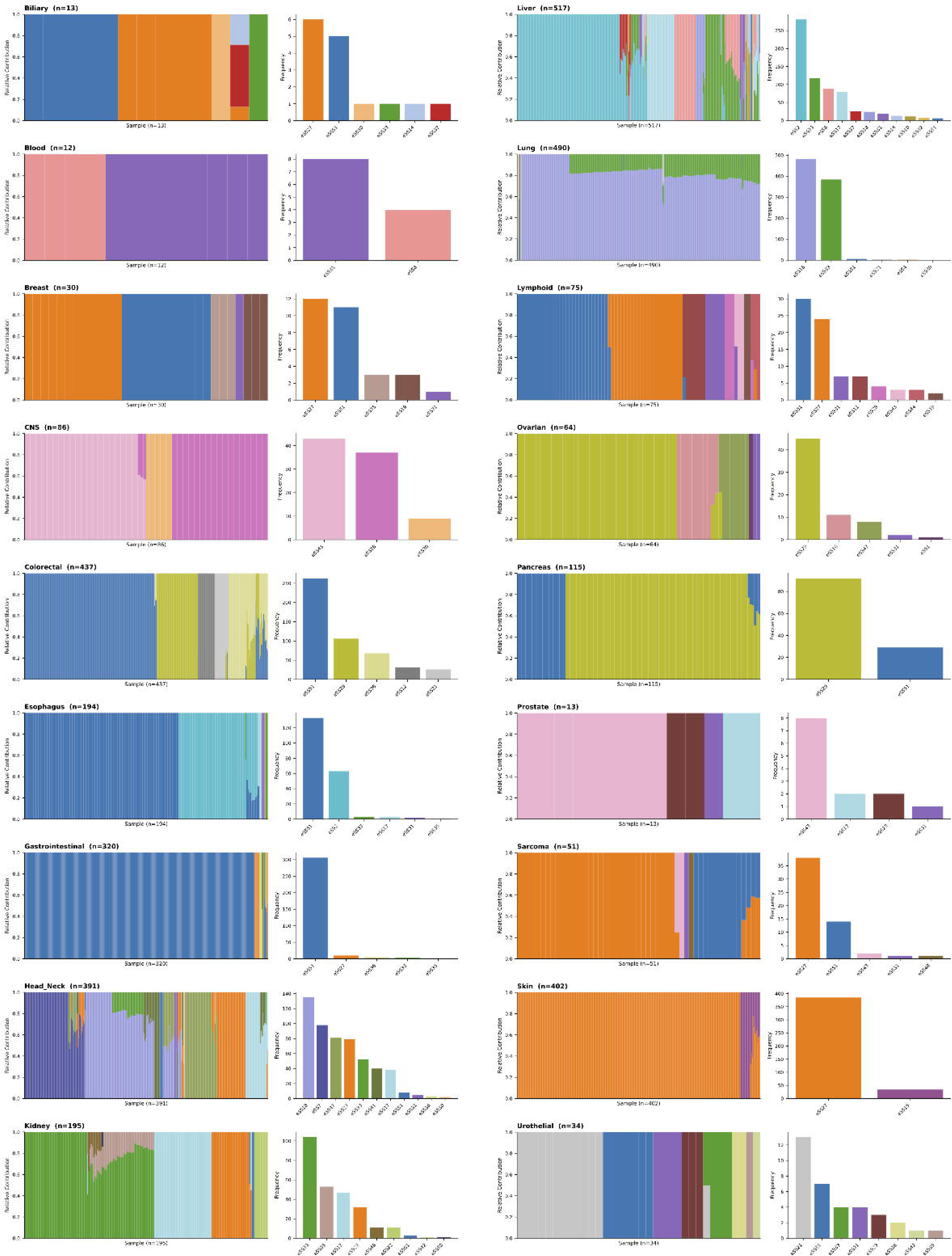
Pan-cancer decomposition overview. Distribution of eSS activity across the 11,428 PCAWG, TCGA, and Mutographs tumors per cancer type.

**Supplementary Figure 17.**
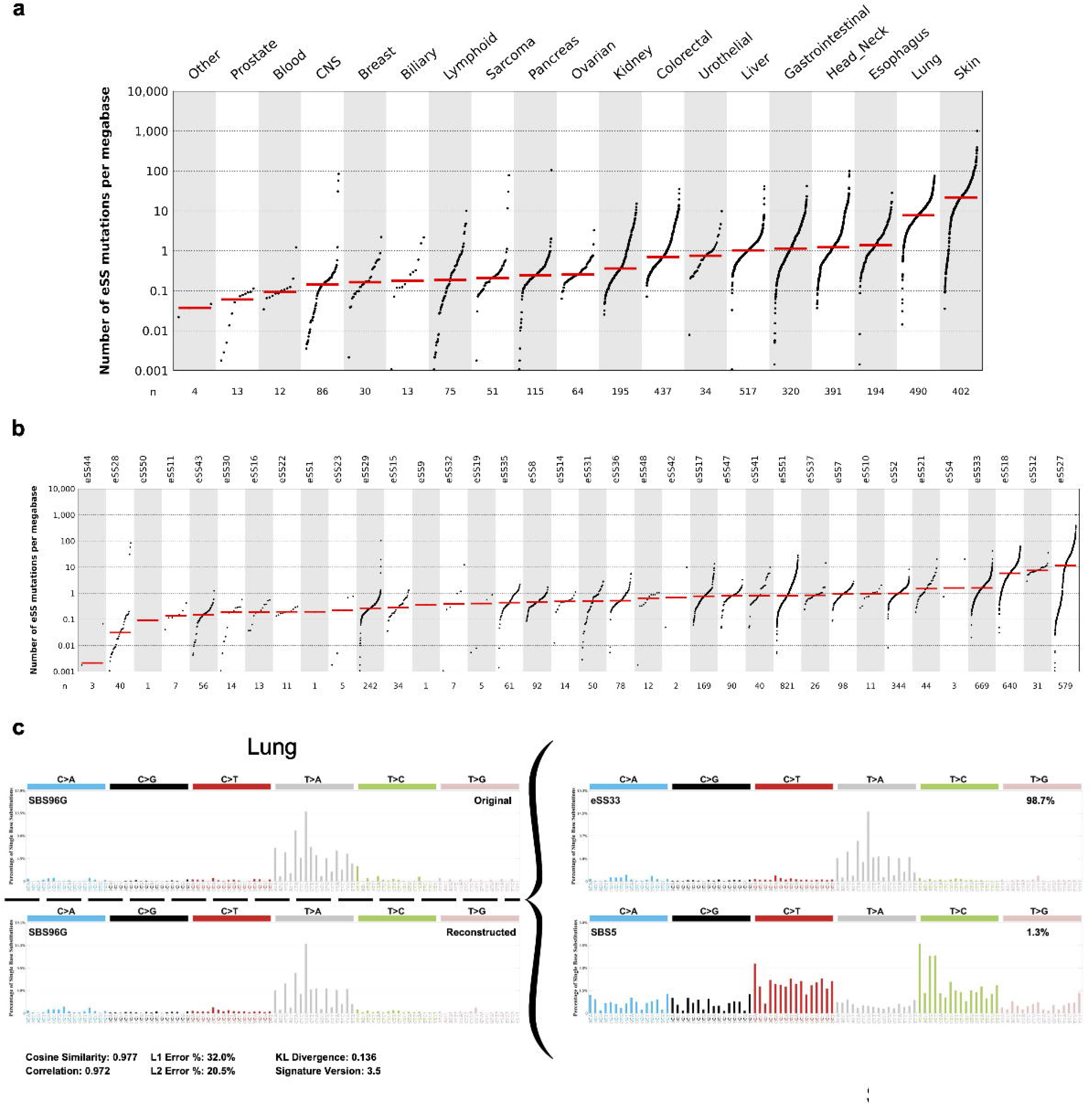
Pan-cancer eSS burden and signature decomposition. (a) Per-sample eSS burden grouped by tissue type. (b) Mutational burden contributed by eSS across pan-cancer cohorts. (c) De novo mutational signature decomposition of SBS96G found in lung cancers into eSS, revealing a strong match with eSS33 that accounts for 98.7% of the profile, with a reconstruction cosine similarity of 0.977.

**Supplementary Figure 18.**
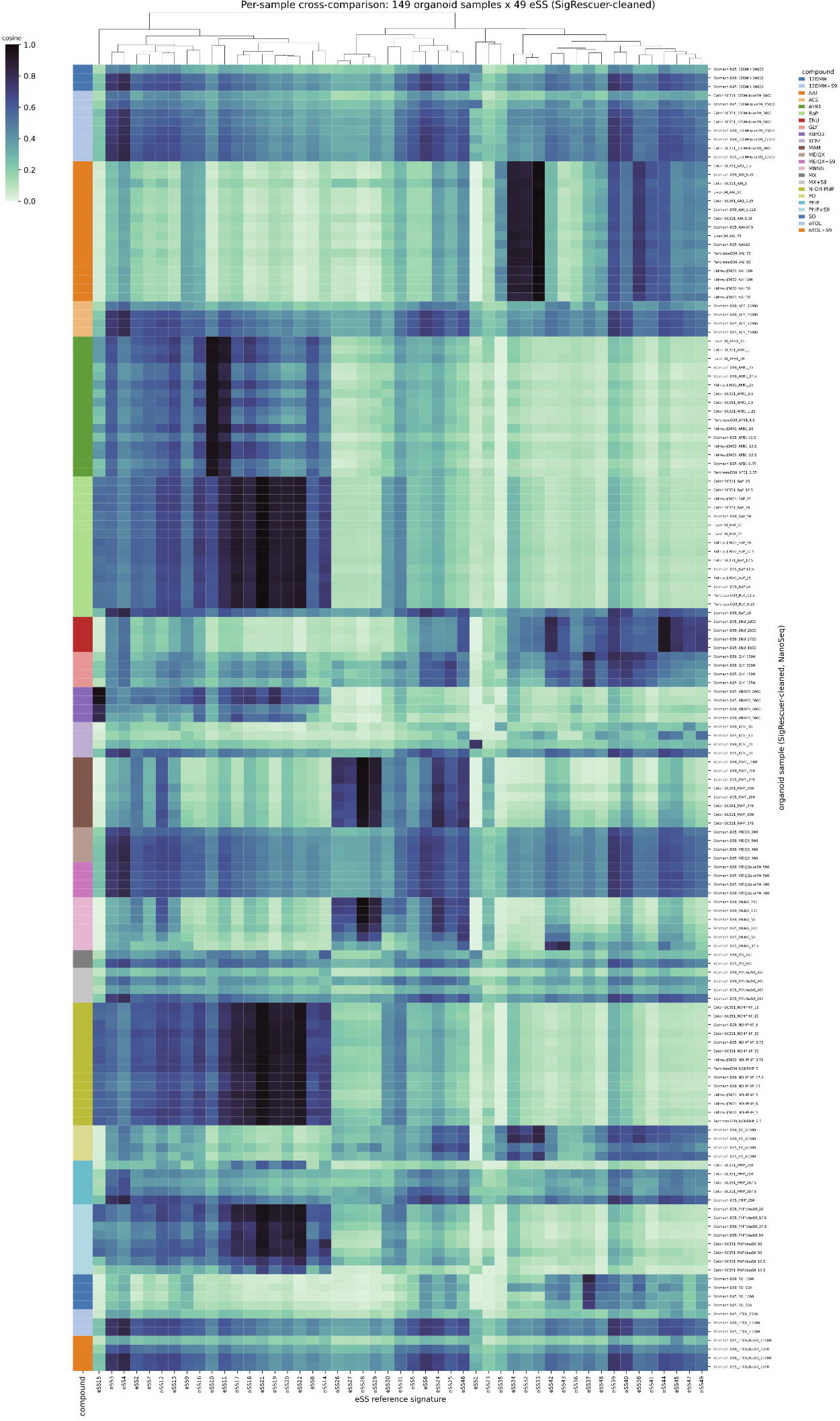
Independent organoid validation by NanoSeq. Cosine similarity of duplex-sequencing organoid signatures against the 49-eSS compendium across tissues, donors, and compounds.

**Supplementary Figure 19.**
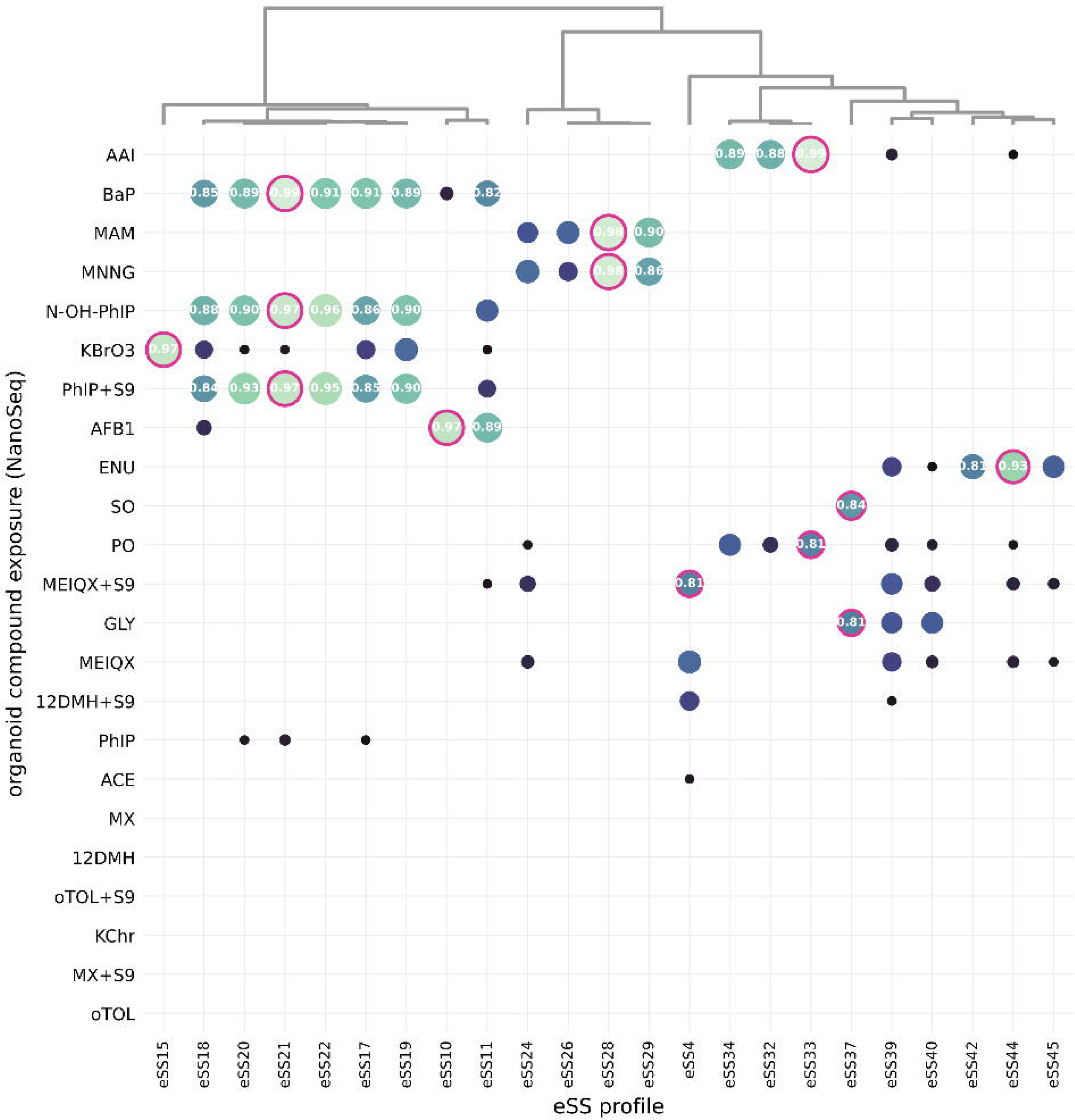
Agent-resolved cosine similarity to the 49-eSS compendium. Clustered matrix of cosine similarity between duplex-sequencing (NanoSeq) organoid mutational profiles, grouped by compound exposure (rows), and the 49 eSS (columns). Dot color encodes cosine similarity (scale 0.6 to 1.0); the best-matching eSS per agent is outlined in magenta and labeled with its cosine value. Compounds without a qualifying match (for example oTOL, MX, ACE, KChr) show no high-cosine cell. The top dendrogram reflects hierarchical clustering of the eSS columns.

**Supplementary Figure 20.**
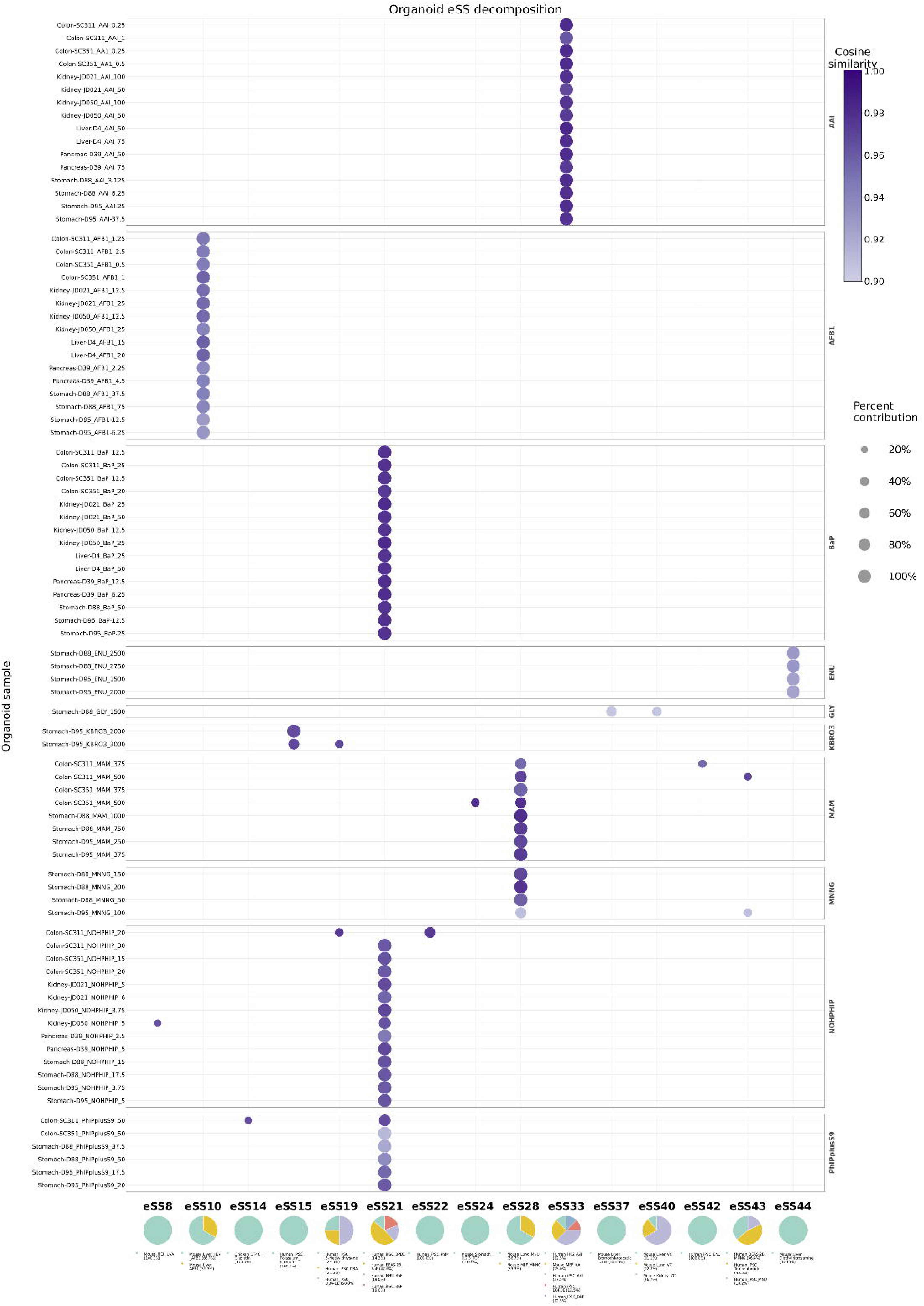
Per-sample eSS decomposition of organoid NanoSeq profiles. Dot plot of eSS assignments across individual organoid samples (rows, grouped by compound) against the eSS atlas (columns). Dot color encodes reconstruction cosine similarity (≥ 0.9) and dot size encodes percent contribution (≥ 5%). Pie charts below summarize the composition of each assigned eSS.

